# Single-molecule imaging reveals a direct role of CTCF’s zinc fingers in SA interaction and cluster-dependent RNA recruitment

**DOI:** 10.1101/2023.12.05.569896

**Authors:** Jonas Huber, Nicoleta-Loredana Tanasie, Sarah Zernia, Johannes Stigler

**Affiliations:** Gene Center Munich, Ludwig-Maximilians-Universität München, Munich, Germany

## Abstract

CTCF is a zinc finger protein associated with transcription regulation that also acts as a barrier factor for topologically associated domains (TADs) generated by cohesin via loop extrusion. These processes require different properties of CTCF-DNA interaction, and it is still unclear how CTCF’s structural features may modulate its diverse roles. Here, we employ single-molecule imaging to study both full-length CTCF and truncation mutants. We show that CTCF enriches at CTCF binding sites (CBSs), displaying a longer lifetime than observed previously. We demonstrate that the zinc finger domains mediate CTCF clustering and that clustering enables RNA recruitment, possibly creating a scaffold for interaction with RNA-binding proteins like cohesin’s subunit SA. We further reveal a direct recruitment and an increase of SA residence time by CTCF bound at CBSs, suggesting that CTCF-SA interactions are crucial for cohesin stability on chromatin at TAD borders. Furthermore, we establish a single-molecule transcription assay and show that although a transcribing polymerase can remove CTCF from CBSs, transcription is impaired. Our study shows that context-dependent nucleic acid binding determines the multifaceted CTCF roles in genome organization and transcription regulation.

## Introduction

Eukaryotic genome architecture, as revealed by high resolution Hi-C maps^1,2^, mirrors the plethora of interactions between different genomic regions. Indeed, two levels of organization have emerged: a global one represented by compartmental domains which are determined by the segregation of transcriptionally active and inactive chromatin regions within the nucleus^3–5^ and a local one where loci of the same domain form strong interactions, leading to the formation of topologically associated domains (TADs)^6,7^. Human CCCTC-binding factor (CTCF) is a transcription factor^8^ that can be found at TAD boundaries^9^. Hi-C experiments showed that removal of CTCF leads to a strong reduction in insulation between TAD domains^10^, indicating that CTCF blocks loop-extruding cohesin complexes^11,12^, and is hence a regulator of chromatin looping.

CTCF consists of 11 zinc fingers (ZFs), that recognize a specific DNA binding sequence (CTCF‐binding site, CBS), and two unstructured termini^13,14^. CBSs are diverse^15^ and share a core sequence that is recognized by the central ZFs of CTCF while upstream motifs are bound by C‐terminal ZFs **(Figure 1A)**^14,16^. CBSs can be further classified into high-and low-affinity binding sites, as certain CBSs have been shown to be more persistent in CTCF depletion experiments^17,18^. Low‐affinity binding sites are often located within genes and at transcription start sites (TSSs), while high-affinity bindings sites can be found at TAD boundaries^19^. This implies that the sequence context modulates CTCF’s activity on different target sites and defines its numerous tasks in genome organization and transcription regulation.

**Figure 1.**
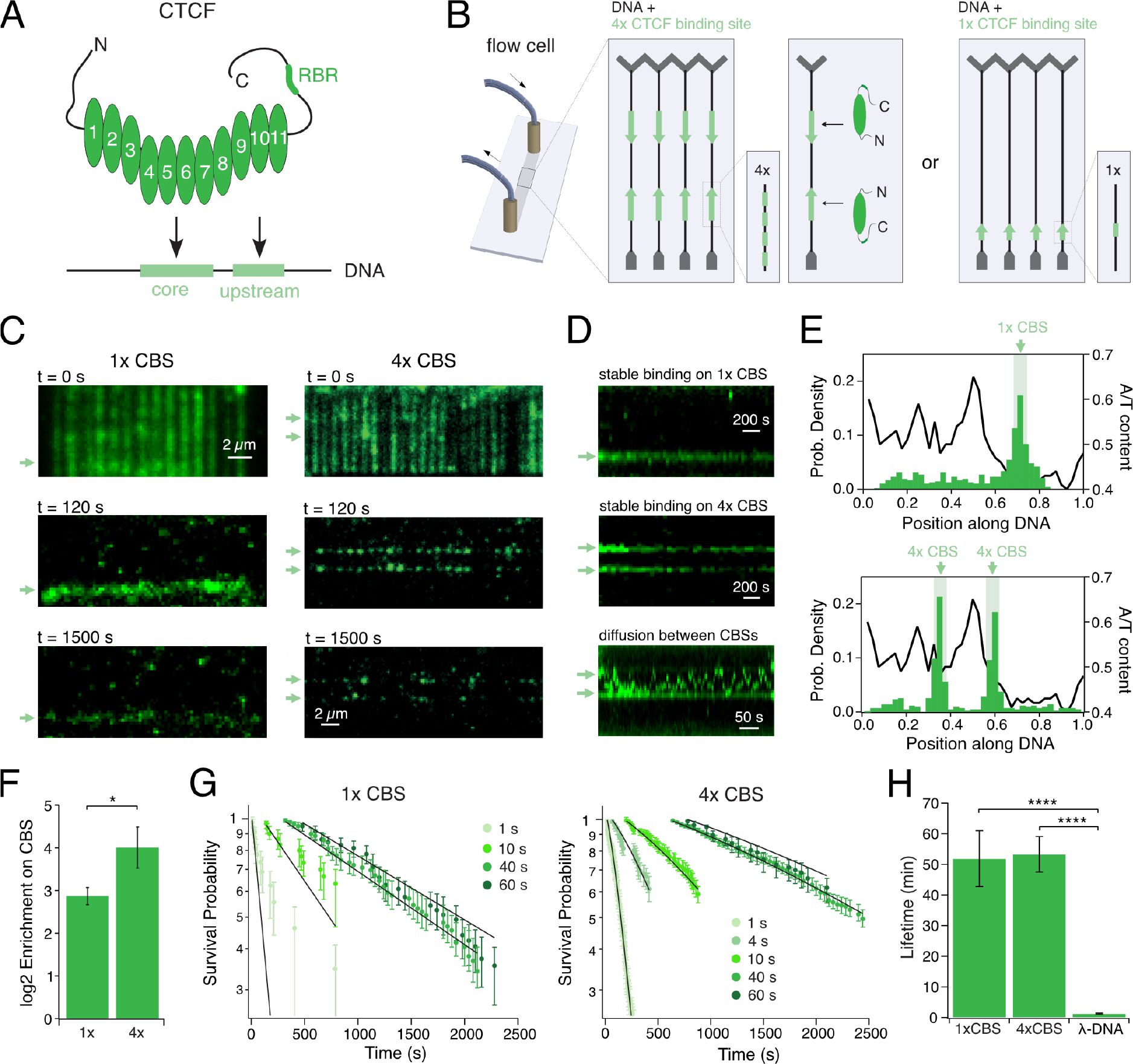
CTCF enriches on both, single and 4x CTCF-binding sites, with a lifetime of ∼50 min. **(A)** Schematic representation of CTCF containing the 11 ZFs, the RNA recognition motif (RBR), the elongated termini and the CTCF binding site (CBS) containing core and upstream motif. **(B)** Schematic representation of the DNA curtains assay. The DNA substrate was designed with either two cassettes of 4x CBSs (light green) with opposing orientation or one cassette with 1x CBS that were included into λ‐DNA (black), which was tethered between Cr barriers (grey) on a custom-built flow cell. **(C)** Representative images of TIRF microscopy of 10 nM CTCF‐WT labeled with Alexa‐Fluor‐568 binding on DNA including either 1x (left) or 4x (right) CTCF binding sites (CBSs, indicated by green arrows). CTCF was first loaded at 50 mM NaCl leading to full coverage of the DNA (top). After washing with 300 mM NaCl, CTCF is enriched on the binding sites (middle) and remains bound for a long time (bottom). **(D)** Representative kymogram of CTCF binding to 1x (top) or 4x (middle) CBSs. Kymogram is shown after a 300 mM NaCl wash. Some non-CBS-bound CTCF diffuse (bottom). **(E)** CTCF enriches at 1x (N = 477) and 4x CBSs (N = 427) after a high salt wash. Light green bars indicate position of CBSs. A/T-content of the DNA substrate is plotted on the right axis. **(F)**CTCF is more strongly enriched (16x) on a 4x than on a 1x site (7x). **(G)**Lifetimes on 1x and 4x CBSs measured at different laser frame rates. A global fit was applied to correct for photobleaching and the presence of multiple CTCFs on the 4x CBSs. **(H)** Photobleaching-corrected CTCF lifetimes on 1x CBS and 4x CBSs are similar, but significantly higher than on λ‐DNA.

A vast majority of CBSs located at TAD borders are convergently oriented^2,20^ i.e., with the CTCF N-termini pointing towards the inside of the TADs. Cohesin is stopped when it encounters the N‐terminus of CTCF. This orientation-dependent arrest of loops is determined by a direct interaction of cohesin subunits SA and Rad21 with the CTCF N-terminus^21,22^. However, *in*‐*vivo* studies revealed that SAs remain associated with CTCF even in the absence of cohesin^23^, suggesting a cohesin-independent interaction between CTCF and SA. Another study showed SA to be the only cohesin subunit directly interacting with CTCF, dependent on CTCF’s C-terminus, in contrast to the Rad21-dependent interaction with its N-terminus^21,24^. Alternatively, the association of CTCF and SA could be facilitated by their shared ability to bind RNA^25–27^. The exact determinants of CTCF-SA interaction remain therefore unknown.

CTCF interacts with RNA through both its outer ZFs ZF1 and ZF10 and an RNA binding domain (RBR) within the disordered C-terminus. This interaction seems to also modulate several aspects of chromatin organization^28,29^. Furthermore, RNA association is linked to CTCF multimerization and cluster formation^26^, which have also been observed in living cells^30,31^. Nevertheless, it remains unclear which CTCF features mediate cluster formation and whether clusters contribute to CTCF’s roles at TADs borders and in transcription regulation.

Complementary to its roles in chromatin looping, CTCF is also associated with gene expression regulation, as it can cooperate with other transcription factors^32^. For example, the TFII-I transcription factor can bind CTCF, and recruit CDK8 for transcription initiation^33^. In addition, CTCF’s role as a transcription regulator is further established by a direct interaction with the largest subunit of RNA Polymerase II (Pol II)^34^. CTCF also acts at later stages of gene regulation as it can influence alternative splicing by promoting Poll II pausing^35^, with the exact mechanism for pausing and recruitment of additional factors thus far undetermined.

CTCF has been attributed many roles in genome regulation, yet mechanistic details of how it fulfills them remain scarce. We set out to ascertain the molecular determinants for CTCF’s various tasks in genome organization and functionality. We purified fluorescently labeled full-length and truncation variants of human CTCF and visualized their association with CBSs using DNA curtains. We show that while a single CTCF is sufficient for target site binding, CTCF’s ZFs mediate clustering on DNA, which enables RNA recruitment. By establishing a single-molecule transcription assay, we show that CTCF is displaced from its CBS by an elongating polymerase, leading to impaired transcription. Furthermore, we uncover a previously unknown mechanism for CTCF association with SA, as we show that SA can be recruited by CTCF bound at CBSs, independently of CTCF termini. Our single-molecule investigation provides insight into how ZF-mediated CTCF interaction with both RNA and DNA modulates CTCF’s various roles at TAD boundaries and in transcription regulation.

## Results

### CTCF has a significantly higher lifetime on CBSs compared to unspecific DNA

CTCF’s target site consists of a core motif bound by ZFs 4‐7 and an upstream motif bound by ZFs 9‐11^14^(**Figure 1A**). CTCF displays increased lifetime on its binding site and off-target diffusion^11,16^. However, it is controversial whether a single CBS is sufficient for CTCF target-site recognition^11,16^. Multiple CBSs create robustness for blocking loop extrusion^36^ and highly conserved topologically associated domains (TADs) in mice contain arrays of CBSs^37^, suggesting that closely spaced CTCF molecules may strengthen TAD borders, perhaps by facilitating protein-protein interactions. Interestingly, CTCF clustering has been shown to occur both *in vivo*^30,31,38^ and *in vitro*^26,38^. To test for CTCF binding cooperativity at the single molecule level, we generated two different λ‐DNA constructs (**Figure 1B**): One construct containing two 4x CBSs in opposite orientation, spaced by 13 kbp resembling a TAD, and the second construct containing only one CBS. End‐modified λ‐DNA molecules were attached to a lipid-bilayer and imaged using TIRF‐microscopy. Fluorescently labeled CTCF was bound to DNA under low-salt conditions, which led to a complete coating of the DNA by the protein (**Figure 1C**, top). Unspecifically bound CTCF molecules were washed off by 300 mM NaCl, while CBS-bound CTCF remained for more than 25 minutes (**Figure 1C**, middle + bottom). Target-site-bound CTCF molecules mostly remained static on the CTCF binding sites even at these high salt concentrations, while single remaining non-CBS-bound CTCF molecules started to rapidly diffuse on DNA (**Figure 1D, Figure S1B**). Photobleaching experiments revealed that neither diffusive nor static CTCF forms higher oligomeric structures (**Figure S1C**). After the 300 mM salt wash, a clear enrichment of CTCF on 1x and 4x CBSs was observed (**Figure 1E**), and enrichment was significantly higher for 4x CBSs (**Figure 1F**). We wondered if this is due to a higher lifetime on 4x CBSs, which would suggest a cooperative binding mechanism. To exactly determine CTCF’s lifetime on its target site and correct for photobleaching we performed measurements at different frame delays for both the 1x and the 4x CBSs (**Figure S1A, Figure 1G**). The data was then analyzed with a global fit model (see methods). For our analysis we assumed three free parameters: the photobleaching lifetime of the fluorescent dye, the lifetime of CTCF on DNA and the chance of each individual site on the 4x CBSs to be occupied with labeled CTCF (see Methods). The global fit resulted in no significant lifetime difference between 1x and 4x sites (**Figure 1H**). CTCF therefore does not bind cooperatively to multiple closely spaced binding sites. The lifetime was more than 40x higher on both 1x and 4x CBSs compared to λ‐DNA.

### CTCF’s inner and outer ZFs but not its termini are required for CBS recognition

We next wanted to identify the roles of the unstructured CTCF termini and the ZFs in CTCF target site recognition. To this end, we produced different truncation mutants missing one (ΔN, ΔC) or both CTCF termini (ΔNC) or some of the ZFs (ΔRBR, ZF9‐CT, ZF4‐7, **Figure 2A**, see supplement for sequences). All mutants containing all 11 ZFs (ΔN, ΔC, ΔNC) were enriched to a similar extent on the CTCF binding sites as the CTCF WT (**Figure 2A and 2B**). In contrast, mutants containing only the upstream-motif-binding ZFs (ZF9‐CT), or the core-motif-binding ZFs (ZF4‐7) were not enriched on the CTCF binding site. Instead, CTCF’s outer ZFs showed a clear preference for AT‐rich regions on λ‐DNA leading to a negative enrichment on the GC-rich CBSs (**Figure 2A and 2B**). Interestingly, a mutant we termed ΔRBR, which lacks the outer ZFs (ZF1, ZF10 and ZF11), that might take part in RNA binding, and the C‐terminal RNA binding domain (RBR)^26,28,29^, was enriched on the CBSs to a similar extent as CTCF WT (**Figure 2B**).

**Figure 2.**
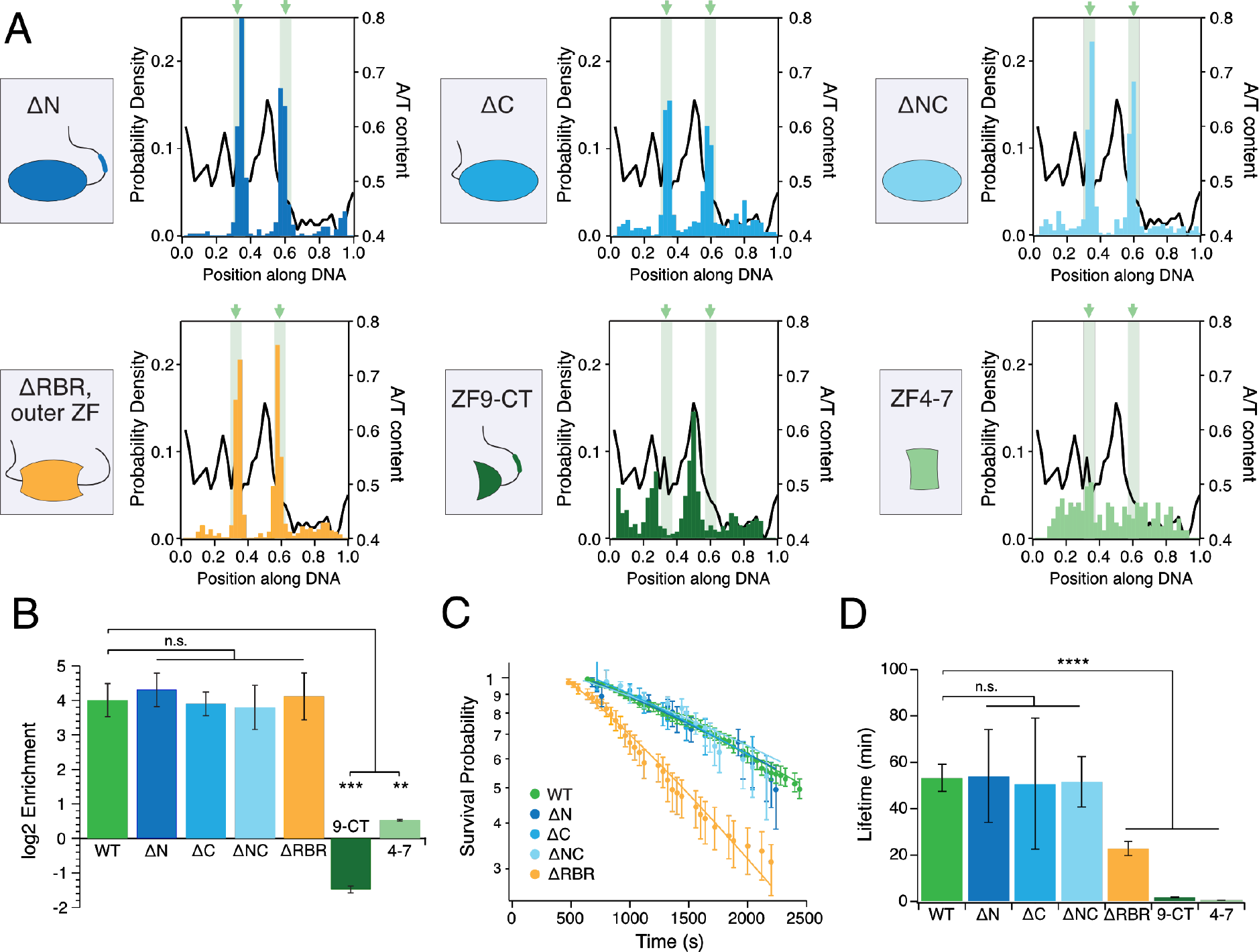
Binding behavior of CTCF variants on DNA curtains **(A)** Binding position of CTCF variants on 4x CTCF binding sites after 300 mM NaCl wash (N = 390/666/396/651/343/88 for ΔN/ΔC/ΔNC/ΔRBR/ZF9‐CT/ZF4‐7). **(B)** Enrichment of CTCF variants on 4x CBSs compared to λ‐DNA. ΔN, ΔC, ΔNC and ΔRBR+ΔZF1,10,11 (ΔRBR) enrich like WT on CBSs. ZF9‐CT (9‐CT) and ZF4‐7 (4‐7) are significantly less enriched than CTCF WT. **(C)** Lifetimes of CTCF variants at 40 s illumination delay. **(D)** Photobleaching-corrected lifetimes of CTCF variants. ΔN, ΔC and ΔNC display similar values like WT. ΔRBR, ZF9‐CT and ZF4‐7 display lower lifetimes.

To test if the different truncation mutations influence CTCF’s stability on DNA, we performed lifetime measurements at different illumination times, similarly to the WT measurements (**Figure 2C and S2A**). Mutants missing one or both of CTCF’s termini but containing all 11 ZF showed no significant difference compared to CTCF WT (**Figure 2D**), while ZF9‐CT had a 25‐fold and ZF4‐7 a 90‐fold reduced lifetime. Interestingly, ΔRBR also showed a 2.5‐fold reduced lifetime, although it was enriched similarly to WT. These data therefore suggest, that the ZFs which recognize the core or upstream motif^16,39,40^ are not stable on the CBS on their own. Instead, binding needs to be stabilized by a combination of core-motif-binding- and outer ZFs. ZFs 9‐11, known to bind to an upstream motif^39^ with a higher AT-content (50 %) compared to the CTCF‐core motif (28 %), thereby preferentially engage AT-rich regions. The presence of AT-rich regions in close proximity to the CBS has been linked to long persistence times of CTCF on chromatin^18^. CTCF missing only some (ZF1, ZF10, ZF11) but not all of these outer ZFs can still recognize its CBS but with a reduced lifetime. Similarly, while ZFs 5‐7 were shown to be essential for binding, the deletion of ZF1, ZF10+11 or ZF9 leads to a significant CTCF half-life reduction on a CBS^41^. In contrast to CTCF’s outer ZFs, CTCF‐termini including the RBR on the C-terminus are not required to increase binding stability of the core-motif-binding ZFs.

### Cohesin’s SA subunits specifically recognize CBS-bound CTCF in absence of other cohesin subunits

After showing enrichment of CTCF on both 1x CBS and 4x CBSs, we asked if CTCF can, according to its role in TADs formation^9^, recruit different parts of cohesin to the CTCF binding site. Most recent data suggest that CTCF blocks loop extrusion via interaction of a conserved domain in CTCF’s N‐terminus with cohesin subunits SA and Rad21^21^. However, CTCF has also been shown *in vivo* to colocalize with SA in the absence of Rad21^23^. The exact mechanism of CTCF’s interaction with SA remains unknown. We sought to test if SA1 or SA2 interact directly with CTCF on the CBSs in the absence of other cohesin subunits. To this end, we purified and fluorescently labeled SA1 and SA2. We then preincubated them with different CTCF constructs before performing a salt wash to remove off-target CTCF (**Figure 3A**). SA1 and SA2 were enriched with CTCF on the CBSs 10‐fold or 8‐fold, respectively (**Figure 3B** and **Figure S3A**), clearly showing that SA subunits can be recruited by CTCF to the CBSs in the absence of other cohesin subunits. Strikingly, even before the salt wash, SA interacted more frequently with CBS-bound CTCF (**Figure 3A**, top). SAs are therefore able to differentiate between CBS-bound and unspecifically bound CTCFs, possibly due to conformation changes in the ZFs when CTCF is able to engage a complete sequence motif.

**Figure 3.**
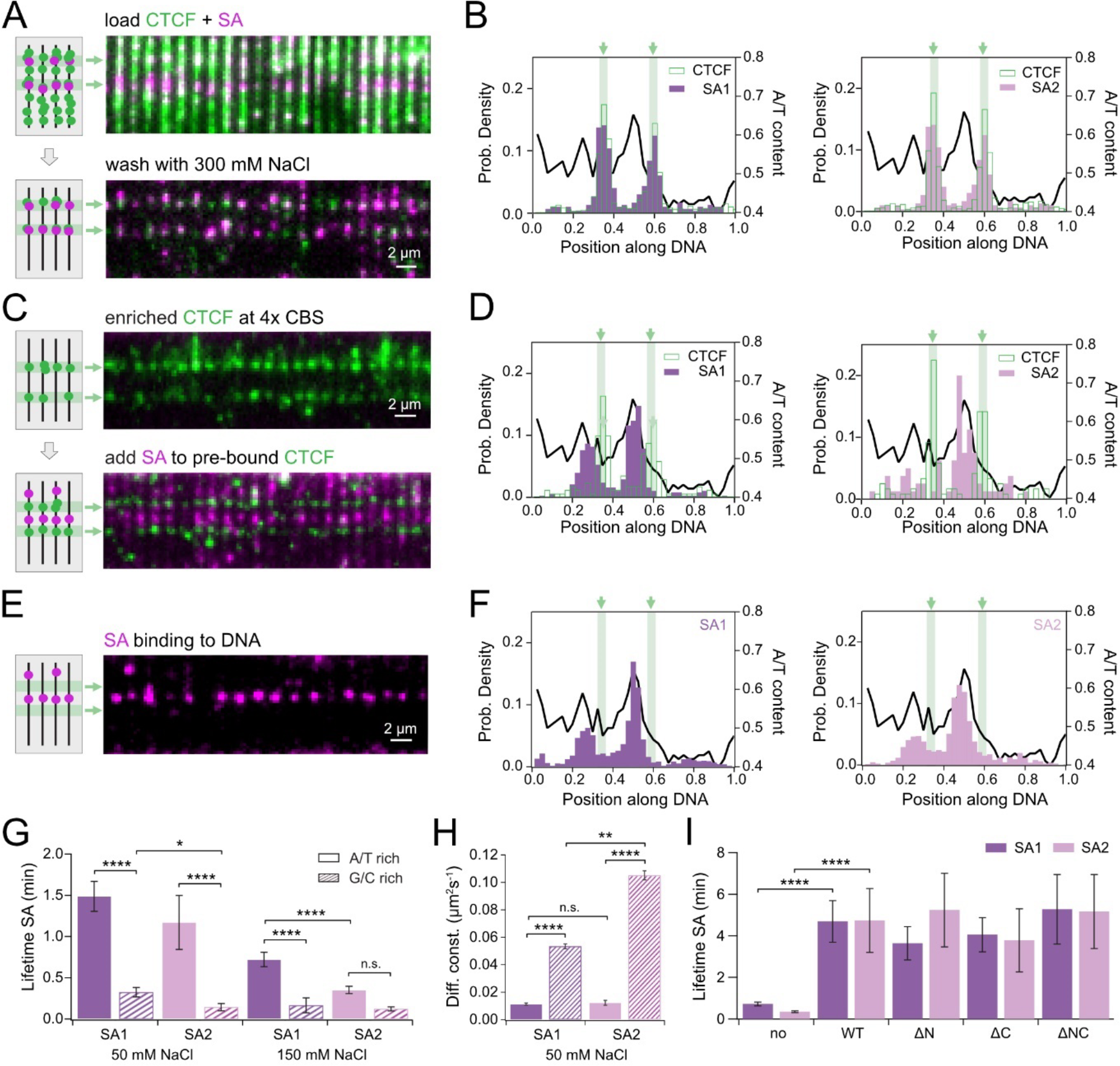
CTCF ZFs stabilize SA on CBSs **(A)** TIRF microscopy of fluorescently labeled CTCF (green) and SA (pink) preincubated before loading on DNA curtains, before (top) and after (bottom) 300 mM NaCl wash. **(B)** Histograms of CTCF (green, left: N = 762; right: N = 747), SA1 (purple, N = 299) and SA2 (pink, N = 196) binding positions after simultaneous load and salt wash. CBSs are shown as green bars and the AT-ratio as a black line. **(C)** TIRF microscopy of CTCF after salt wash (top) followed by SA load (bottom). Color code as in (A). **(D)** Histograms of CTCF (green, left: N = 171; right: N = 120), SA1 (N = 380) and SA2 (N = 80) binding positions after sequential load. **(E)** TIRF microscopy of SA (magenta) binding on DNA‐curtains. **(F)** Histograms of SA1 (N = 1321) and SA2 (N = 875) binding positions on λ‐DNA. **(G)** Lifetime of SAs on AT-rich and GC-rich DNA regions at 50 mM and 150 mM NaCl. At 50 mM NaCl, both SAs have a higher lifetime on AT-rich than on GC-rich regions. At 150 mM, only SA1 shows a higher lifetime on AT-rich, which is also significantly higher than for SA2. **(H)** SA1 and SA2 diffusion coefficients on AT-rich and GC‐rich regions (coloring like in G). SA1 and SA2 diffuse significantly faster on GC‐rich than on AT‐rich regions. SA2 diffuses significantly faster than SA1 on GC‐rich regions, but not on AT‐rich regions. **(I)** Lifetime of SAs in absence of CTCF and after recruitment to CBSs by different CTCF constructs. SA1 and SA2 have a significantly higher lifetime on CTCF WT than on DNA. Both SAs have similar lifetimes on CTCF variants compared to CTCF WT.

### SAs target AT-rich regions on DNA

To directly observe if SA can specifically recognize CTCF once it has already bound its CBS, we first enriched CTCF on the CBSs and then loaded SA1 or SA2 (**Figure 3C**). Surprisingly, this led to a completely different binding behavior. Instead of colocalizing with CTCF, both SA1 and SA2 showed a clear preference for AT-rich DNA regions (AT-content ≥ 50 %) (**Figure 3D**). A possible explanation could be that pre-bound CTCF blocks GC-rich regions. We therefore tested the binding behavior of SA without CTCF and again found a clear preference for AT-rich regions (**Figure 3E, F**). By comparing lifetimes **(Figure 3G)** and diffusion **(Figure 3H, Figure S3B)** at low salt on AT-rich (AT content ≥ 50 %) and GC-rich (< 50 %) regions, we found this enrichment to happen by faster diffusion on GC-rich regions and by a lower lifetime on GC-rich regions leading to more rapid sampling. At higher salt concentration (150 mM NaCl), SA quickly unbound from DNA (**Figure 3G**). SA diffusion was also observed on DNA previously enriched with CTCF (**Figure S3C**). When SA collided with a CBS-bound CTCF, we observed a large fraction of intermediate and long-term binding events (>20 sec, **Figure S3D**). Less frequently, SA was blocked by CTCF and changed its diffusion direction. In very rare cases, we observed SA passing a CBS-bound CTCF (**Figure S3C** and **S3D**). We conclude that diffusing SAs can recognize CBS‐bound CTCFs on DNA. However, since our DNA substrate contains large AT-rich regions, where SA accumulates, this recruitment is comparatively rare.

### SAs interact more stably with CTCF’s ZFs than with DNA

Since diffusion to the CBSs seems to be an inefficient mechanism of SA recruitment, we asked how SA is enriched on the CBSs in our combined loading experiments. In contrast to the low salt stability of SA on DNA, SA preincubated with CTCF and recruited to the CBSs stayed bound for multiple minutes at 300 mM NaCl **(Figure 3I)**. This argues that CBS-associated SAs are not bound to the DNA but directly attached to CTCF.

We next sought to find out if this interaction is further stabilized by cohesin’s Rad21 subunit. We therefore copurified SA1 or SA2 with a Rad21 peptide containing its CTCF binding region^21^ and validated its interaction with SA by mass photometry (**Figure S3E**). Rad21 did not influence SA enrichment on CTCF in sequential or simultaneous load experiments and had no influence on SA lifetimes on CTCF (**Figure S3F–I**). Rad21 therefore does not seem to be required for the CTCF‐SA interaction.

It is known that a conserved region in the N‐terminus of CTCF is required for cohesin interaction^21,29,42^. To test whether this is also true for the interaction with SA, we incubated CTCF truncation variants with SA1 or SA2 and loaded them on DNA curtains. No significant differences in interaction times **(Figure 3G)** or enrichment (**Figure S3A**) were observed for any truncation mutant. We therefore conclude that SA is recruited to and stabilized on CBSs by specifically binding to CBS-bound CTCF via a ZF-mediated interaction.

### CTCF and CTCF-SA complexes do not block transcription

CTCF has a well-established role as a transcriptional insulator, blocking distant enhancer-promoter interactions in vertebrates^43,44^. CTCF might also directly influence transcription by interaction with the RNA‐Polymerase II via its C‐terminal domain^34^. Binding of CTCF near TSSs can influence Pol II processivity *in vivo*, which impacts mRNA splicing efficiency^35,45–47^. Our goal was to test both the influence of CBS-bound CTCF and CTCF-SA complexes on transcription and the influence of a transcribing polymerase on these complexes. Hence, we performed *in*‐*vitro* transcription assays both in bulk and single-molecule using T7‐RNA‐polymerase (T7‐Pol). For bulk measurements, we used two linear constructs containing construct-centered 4x CBSs in two orientations downstream of a T7‐promoter (**Figure 4A,B**). Unexpectedly, neither the amount nor the length of produced RNA was influenced by CTCF on both type of templates, suggesting that CTCF does not inhibit transcription (**Fig. 4A,B**). CTCF was therefore either passed, pushed or removed from the DNA by the transcribing polymerase.

**Figure 4.**
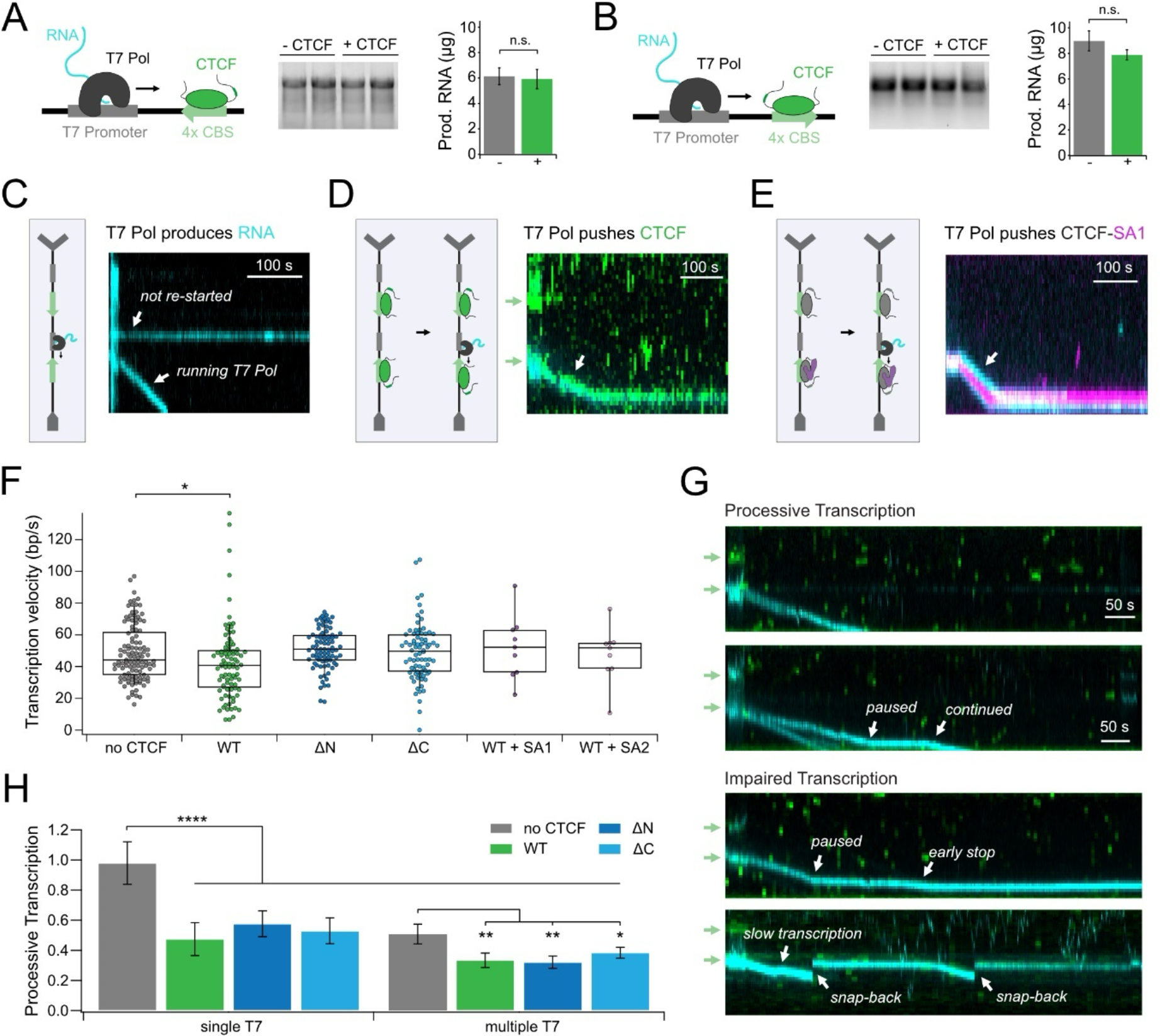
CTCF does not block transcription by T7 polymerase. **(A)** T7‐Pol bulk transcription assay on a PCR product containing a T7‐promoter as well as 4x CBSs with the downstream motif pointing towards the promoter. The length and amount of produced RNA was measured for three independent experiments with and without CTCF and no significant reduction in amount of produced RNA was observed. **(B)** Same as (A) but with upstream motif pointing towards the promoter. **(C)** Illustration of in‐vitro transcription assays and representative kymograms for T7‐Pol moving down the DNA during transcription (cyan = Cy3‐labeled RNA). **(D)** same as (C) but after enrichment of CTCF on 4x CBSs. CTCF (green) and T7‐Pol (cyan, RNA) are moving mutually. **(E)** same as (C) but after enrichment of SA-CTCF complexes containing unlabeled CTCF and LD555-labeled SA (magenta) on 4x CBSs. CTCF-SA complexes are pushed by T7‐Pol. **(F)** Mean transcription velocities of T7‐Pol alone and T7‐Pol pushing different CTCF variants or CTCF-SA complexes. WT CTCF reduced transcription velocity, while no significant difference was found for ΔN, ΔC, SA1-CTCF and SA2-CTCF compared to T7‐Pol velocities. **(G)** Representative kymograms of continuous transcription (top), pausing and stopping events (middle), and snapback of polymerases (bottom). **(H)** Fraction of processive transcription in case of single or multiple transcription events on one λ‐DNA molecule. All CTCF variants impair transcription significantly in both cases of single and multiple T7 polymerases on one DNA. In absence of CTCF, multiple T7‐Pols on one DNA impair transcription also significantly.

To test these possibilities, we performed single-molecule transcription assays. For this, we altered our previously used λ‐DNA‐construct by inserting a T7‐promoter in front of each 4x CBSs (**Figure 4C**). Transcription assays were then performed in two steps. First, transcription was carried out for 2 minutes in the presence of labeled nucleotides (Cy3‐UTP) to produce fluorescently labeled RNA. Second, after a brief wash to remove the labeled nucleotides, transcription was restarted in the presence of only dark nucleotides to reduce the fluorescent background in the flowcell. The movement of the transcribing T7‐Pol was followed by tracking the Cy3-labeled RNA (**Figure 4C**).

To test CTCF’s influence on transcription we loaded and enriched fluorescently labeled CTCF on the CBSs as described. After restart, transcribing T7‐Pols pushed CTCF off its binding site. CTCF (green) and RNA (cyan) fluorescence co-localized during the whole transcription process (**Figure 4D**). We compared the velocities of these pushed complexes to those of T7‐Pol alone and found a small but significant reduction in transcription velocity in the presence of CTCF WT (p= 0.034) (**Figure 4F**). In contrast, the ΔN and ΔC variants did not slow down T7‐Pol. However, consistent with bulk data (see above), these differences were small. We therefore conclude that CTCF does not block transcription.

Our single molecule data revealed a stable interaction between CBS-bound CTCF and SA (see above). We therefore wondered whether SA-CTCF complexes show a different behavior than CTCF alone when encountered by a transcribing polymerase. However, SA-CTCF complexes (SA labeled, CTCF dark) were also pushed by T7‐Pol (**Figure 4E**). We conclude that neither CTCF alone nor SA1-CTCF nor SA2-CTCF complexes block transcription.

### CTCF increases the frequency of impaired transcription events

In the absence of CTCF, most T7‐Pol transcription elongation events (98 %) were processive (**Figure 4H**). Strikingly, in the presence of CTCF WT, ΔN or ΔC, we observed a significant reduction of processive transcription to 48-58 % and an increased amount of permanent early stopping and snap-backs (**Figure 4G**). After being pushed off its CBS, CTCF might get evicted from DNA by the nascent RNA chain. CTCF-RNA binding has been linked to the formation of transcription clusters *in vivo*^31^. We therefore hypothesized that impaired transcription in our experiment is a consequence of increased T7‐Pol collision events in CTCF-RNA clusters. To test if clustered or closely spaced polymerases impair transcription^48^, we split our data into DNA molecules on which we only observed a single transcription event, versus multiple transcription events. Interestingly, even in the absence of CTCF, multiple transcribing polymerases on the same DNA led to a significant decrease in processive transcription to 51 %, which is even further reduced in presence of CTCF to 32-38 % **(Figure 4H)**. We thus propose that CTCF increases interactions between polymerases by forming local clusters with RNA^26^, which then disturbs transcription.

### CTCF oligomerization is required for secondary RNA-capture

CTCF increases the frequency of polymerase stalling and reduces the fraction of processive polymerases (see above). We therefore asked if this processivity reduction stems from DNA-bound CTCF binding to the nascent RNA, i.e., secondary RNA capture. To test this possibility, we enriched CTCF on 4x CBSs as shown above (**Figure 1C,E,F**) and incubated with fluorescently tagged RNA in solution (**Figure 5A**). Unexpectedly, we observed no RNA recruitment to CBS-bound CTCF. CTCF-RNA binding has been linked to CTCF clustering^26,28^. However, whether RNA binding is required for oligomerization or *vice versa* remains elusive. To test if CBS-bound CTCF is monomeric, we performed photobleaching experiments (**Figure 5B**). We found that most fluorescent puncta at 4x CBSs bleached in less than four steps, with an average of 2.5 ± 0.1 steps (**Figure 5C**). At a pre-determined labeling efficiency of 52 ± 8 %, this is consistent with monomeric CTCF. CTCF cluster formation might require a high local concentration of DNA-bound CTCF^38^. We therefore repeated the RNA capture experiments before removing unspecifically bound CTCF. In contrast to our previous result, RNA was efficiently captured and even remained bound after a high-salt wash (**Figure 5D**). CTCF photobleaching analysis at positions of captured RNA revealed higher oligomeric structures with an average of 8.8 ± 0.3 steps, corresponding to an average of 17 CTCF molecules (**Figure 5E,F**).

**Figure 5.**
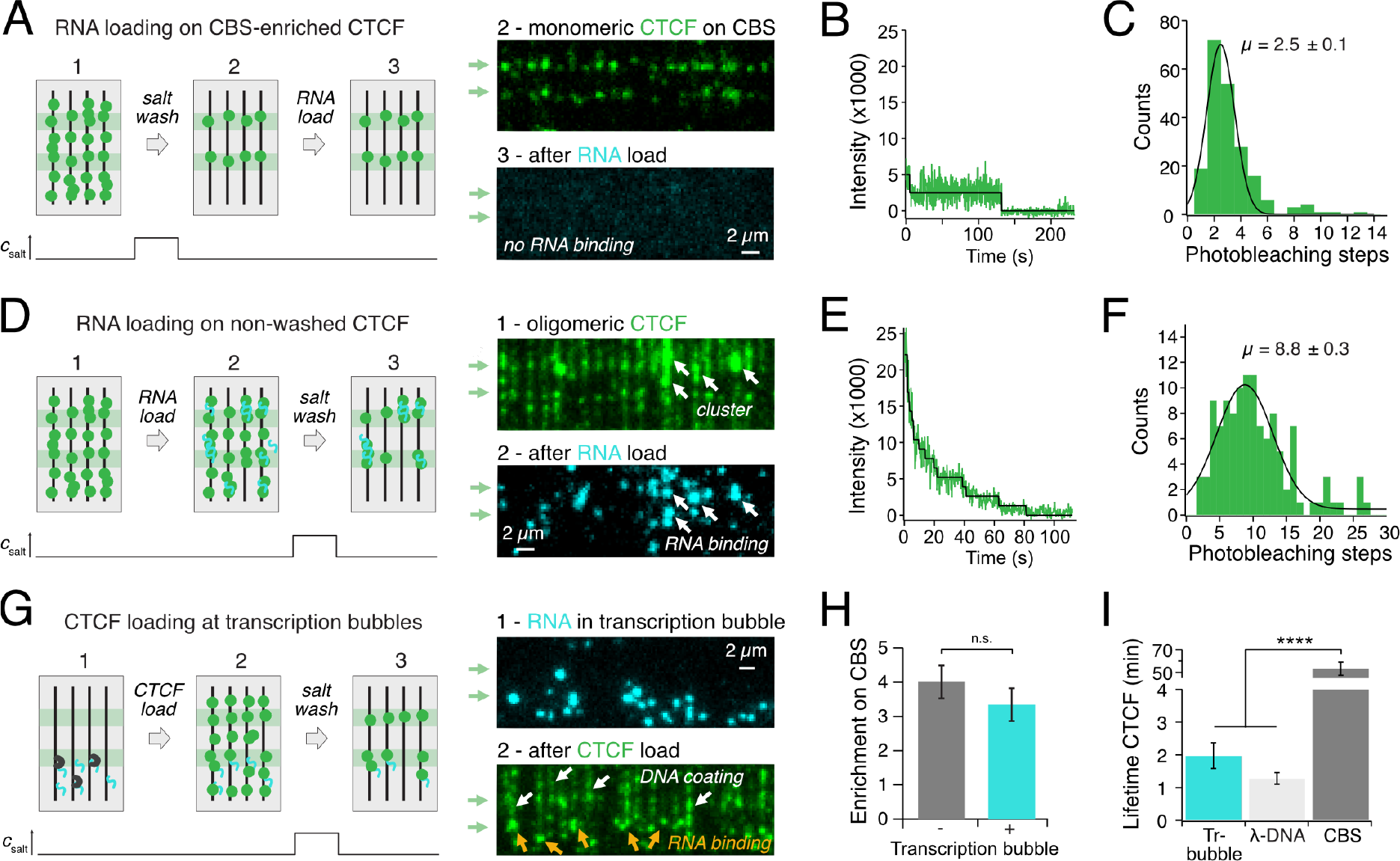
RNA recruitment by oligomeric CTCF. **(A)** RNA-loading on CBS‐enriched CTCF. Left: Experimental workflow. Right: TIRF microscopy of CBS‐bound CTCF (green) before RNA load (top) and of Cy3-labeled RNA (cyan) after RNA load (bottom). No RNA capture was observed. **(B)** Representative intensity curve of a two-step CTCF photobleaching event. Bleaching steps are illustrated by a black line. **(C)** Histogram of CTCF bleaching steps on 4x CBSs. Same as Figure S1C, added here for comparison. **(D)** RNA loading on clustered CTCF. Left: Experimental workflow. Right: TIRF microscopy of CTCF clusters (green) before RNA load (top) and of Cy3-labeled RNA (cyan) after RNA load. RNA is recruited to CTCF clusters (white arrows). **(E)** Representative intensity curve of multi-step CTCF bleaching event of a CTCF-RNA cluster. Steps are illustrated by a black line. **(F)** Histogram of CTCF bleaching steps in CTCF-RNA clusters. **(G)** CTCF loading at transcription bubbles. Left: Experimental workflow. Transcription bubbles were formed by loading T7‐Pol and facilitating transcription as before. Right: TIRF microscopy of Cy3-labeled RNA in transcription bubbles (cyan) before CTCF load (top) and of CTCF (green) after loading (bottom). CTCF is partially recognizing transcription bubbles (orange arrows), but is mainly coating the DNA (white arrows). **(H)** Enrichment of CTCF on 4x CBSs is not significantly different in presence or absence of transcription bubbles. **(I)** CTCF lifetime on transcription bubbles is similar to lifetimes on λ‐DNA and significantly smaller than on 4x CBSs.

To analyze the role of the CTCF-termini in multimerization and RNA-capture, we repeated the same experiments for the CTCF ΔNC mutant. Interestingly, the mutant still formed multimers on DNA (9.0 ± 0.5 bleaching steps), which in turn were able to capture RNA (**Figure S4D**). Hence, the termini of CTCF are not required for RNA capture and oligomerization is caused by ZF interactions^16,38^. On the contrary, CTCF’s termini might even impede cluster formation to some extent since the bleaching step histogram of CBS-enriched ΔNC displays a second peak not observed for any other mutant **(Figure S2B)**.

WT- and ΔNC-RNA clusters were enriched 4‐fold on the 4x CBSs, but this enrichment was significantly smaller than for monomeric CTCF without RNA (**Figure S4A**,**C**): More CTCF-RNA clusters stayed bound at unspecific positions, implying that RNA binding additionally stabilizes CTCF on low-affinity binding sites. In contrast to CTCF, DNA-bound SAs did not capture RNA (**Figure S4E**). Instead, both SA1 and SA2 were mostly washed off the DNA by RNA, indicating a higher affinity of SAs for RNA than DNA (**Figure S4F**)^25^. We conclude that while RNA binding removes SA, it stabilizes clustered CTCF and does not bind to monomeric CTCF.

### Both CTCF and SA colocalize with transcription bubbles

Since CTCF is involved in the formation of transcriptional condensates^31^, we wanted to test if CTCF’s ability to bind RNA causes direct recruitment by transcription bubbles. We performed transcription experiments like described above, followed by CTCF incubation. We found that CTCF colocalized with transcription bubbles (**Figure 5G, Figure S4B**), but there was no significant decrease in CBS enrichment, as CTCF prefers CBSs over RNA (**Figure 5H**). Lifetime on CBSs was more than 25 times larger than on RNA **(Figure 5I)**.

In contrast, when we repeated this experiment with SAs, which colocalize with R‐loops *in vivo*^23^, the presence of transcription bubbles on DNA curtains led to a complete change in binding behavior (**Figure S4G**,**H**). Both SA1 and SA2 were no longer enriched on AT-rich regions (**Figure S4I**) and instead accumulated at transcription bubbles, with a higher lifetime compared to DNA (**Figure S4J**). This is consistent with our previous observation of SA preferring RNA over DNA (**Figure S4F**). We conclude that both CTCF and SA can be recruited by RNA transcription bubbles. However, while this recruitment drastically changes the DNA localization of SA, CTCF is still mostly localized at CBSs.

## Discussion

### CTCF’s DNA binding dynamics are influenced by RNA and transcribing polymerases

CTCF influences a wide variety of processes in mammalian cells by binding to its genomic target site, including insulation^43,49,50^, alternative splicing^35,46,51^, transcription activation^52^ and TAD boundary formation^6,7,9^. Our study showed that CTCF binds stably on CBSs, while it displays diffusion and low lifetimes on unspecific DNA **(Figure 1C-E)**^11,12^. By performing measurements at different laser frame rates and correcting photobleaching data, we showed that CTCF has a much higher lifetime on CBSs than previously observed *in vitro*^11,12^ and also *in vivo*^39,53–57^. The lower lifetime *in vivo* might result from interactions with other chromatin bound proteins, as CTCF in resting B‐cells showed a lifetime similar to our results^55^. In our transcription assay, CTCF and CTCF-SA were pushed off CBSs by a transcribing polymerase **(Figure 4D-F)**, which supports the theory of chromatin-bound proteins reducing CTCF’s lifetime on DNA, an effect which might be weaker in resting B‐cells, where transcription elongation is repressed^55^. Our data is therefore consistent with a model where transcription impacts CTCF’s residence time on chromatin. Transcription can displace cohesin *in vitro*^58^, disrupt its localization at CTCF sites^59^ and act as a mobile loop extrusion barrier^60^. CTCF anchors near active genes might therefore be less stringently positioned and cause more diffuse interactions^61^.

Correspondingly, we also observed CTCF to have an impact on transcription. Single-molecule transcription assays revealed a higher amount of impaired transcription and a slower transcription rate in the presence of CTCF **(Figure 4F-H)**. A study on a bacterial RNA‐Pol has shown that the length of transcription pauses, and the appearance of backtracking events is increased by transcription-opposing forces created by the nascent RNA^62^. We speculate that CTCF, once pushed off its CBS by the polymerase, might be evicted from DNA by the nascent RNA^63,64^ and change the secondary structure of the transcript. Permanently paused states could also be caused by polymerase unbinding and CTCF capturing the RNA chain. Transcription could therefore enable CTCF to recruit RNA to CBSs and lead to an increased insulation at domain boundaries^65^. Impaired transcription by CTCF could also regulate Pol II pausing^47^, which has been linked to alternative splicing^35^.

A different explanation for transcription impairment is CTCF clustering and consequential DNA or RNA crosslinking^26,38,66^ causing an increased number of T7 collisions, which can lead to T7 displacement from DNA^48^. CTCF clustering has been linked to RNA binding^26,28^, but it remains elusive whether CTCF’s interaction with RNA leads to the formation of clusters or if clustering is required for RNA capture. Here, we show that DNA-bound CTCF forms clusters, and that only clustered CTCF can capture RNA **(Figure 5A-F)**. Oligomerization therefore enables RNA recruitment by chromatin-bound CTCF. Photobleaching experiments revealed an average number of 17 CTCF molecules per cluster, which is close to values observed in the nucleus^30^. However, cluster sizes might differ in living cells, as both cohesin and transcription can shape cluster formation, with the former facilitating and the latter disrupting it^30^.

Our data show that RNA modulates CTCF binding positions and stability *in vitro*. In the absence of RNA, CTCF is only stably bound to high-affinity binding sites, while RNA binding leads to stable binding also at low-affinity sites (**Figure S4A**,**C**). Correspondingly, *in vivo*, low-affinity CBSs flank regions of active chromatin and have been associated with transcriptional regulation, while high-affinity CBSs more often flank repressed chromatin and are associated with regulating genome architecture^19^. CTCF on high-affinity CBSs is highly stable^17,18^, and loop domains can persist for hours without energy input, requiring stable anchoring by CTCF^67^, in agreement with our observed long lifetimes on CBSs. Binding to low-affinity binding sites is less persistent^19^ and disrupted by transcription inhibition or RNA-binding deficient mutants^29^. Our results therefore support a model where RNA transcripts are an important regulator of CTCF binding positions and stability. This means that RNA is a key regulator in CTCF function, allowing a dynamic modulation of CTCF binding near active genes, but is less critical for CTCF’s role in genome architecture. The latter is facilitated by CTCF’s high CBS-stability, independently of clustering and RNA binding.

Our data demonstrate that oligomerization and RNA capture is independent of the unstructured termini **(Figure S4D)**. On the contrary, the termini might even impede cluster formation **(Figure S2B)**. CTCF is assumed to bind to RNA via its ZFs 1 and 10 as well as a C‐terminal RNA binding region^28,29^. However, a more recent study shows a ncRNA to interact with ZF3‐6 and impede CTCF binding to its genomic locus^68^. Since CTCF can bind RNA with multiple ZFs, and we found no RNA capture for monomeric CBS-bound CTCF, we propose a model in which CTCF oligomerization creates unoccupied ZFs, enabling simultaneous RNA and DNA-binding. This would allow CTCF to create an RNA interaction hub on chromatin possibly involved in the formation of transcriptional condensates^31^ and RNA-dependent recruitment of cohesin^69^.

### Both CTCF and RNA can stabilize SA-DNA binding independent of cohesin

SA colocalizes with R‐loops *in vivo*, possibly by a direct interaction with RNA, or with other RNA-binding proteins^23^. Our experiments show that in the absence of RNA, SA preferentially interacts with AT-rich DNA-regions, with below-average occupancy on CBSs, fast binding dynamics and low salt stability **(Figure 3F-I)**. In contrast, SA displayed stronger colocalization with and a significantly more stable binding at transcription bubbles than on DNA. **(Figure S4G-J)**. SA-RNA interactions could therefore facilitate their localization to R‐loops and to active genes^23,70–72^, as well as recruitment of cohesin‐SA to CTCF sites^69^. However, unlike CTCF, DNA-bound SA cannot recruit RNA. Instead SA is displaced from DNA by RNA **(Figure S4E-F)**, arguing that SA’s binding site for RNA and DNA are identical, with higher affinity **(Figure S4J)**^25^ for RNA over DNA.

CTCF can directly recruit SA independently of RNA and other cohesin subunits, by significantly increasing lifetime and salt stability compared to DNA-bound SA **(Figure 3I)**. Moreover, the CTCF termini are not required for interaction with SA, suggesting a ZF-mediated recruitment mechanism of SA to CBSs **(Figure S3A)**. SA enrichment is enhanced as it seems to be able to specifically recognize CTCF on CBSs **(Figure 3A)**, most likely through recognizing different sequence-induced CTCF binding modes. CTCF is more salt-stable than any cohesin subunit on chromatin^73^, and CTCF depletion leads to a decrease in cohesin residence time^74^. We therefore propose that direct SA-ZF interactions increase the residence time of the complete cohesin‐complex at TAD‐borders, enabling the formation of long-lived loops^67^. Two genome localization studies revealed distinct subpopulations of SA1‐cohesin and SA2‐cohesin^75,76^. Interestingly the SA2-cohesin subpopulation displayed a comparatively lower colocalization with CTCF. Here we show that SA in the absence of CTCF is not stably associated to DNA at physiological salt concentrations **(Figure 3G,I)**, in line with the SA2‐cohesin subpopulation displaying a low residence time^76^ and salt stability^75^. In contrast, we show that SA-ZF interactions stabilize SA even above physiological salt concentrations on CBSs **(Figure 3I)**. This could explain why the SA1‐cohesin subpopulation in these studies, colocalized with CTCF at TAD borders, displayed a higher salt stability^75^ and residence time^76^. Our data therefore suggest that CTCF-mediated SA stabilization on CBSs can mediate different binding dynamics and roles of SA1‐cohesin and SA2‐cohesin^71,72,75^. However, as we observe a similar stabilization of SA1 and SA2, it remains to be determined what other factors regulate the increased colocalization of CTCF with SA1‐cohesin compared to SA2‐cohesin *in vivo*.

Cohesin’s Rad21 subunit was previously shown to be essential for the interaction between SA and the CTCF N‐terminal SA recognition motif^21^. However, another study showed that Rad21 is dispensable for an interaction between SA and the CTCF C‐terminus^24^. In our experiments, we did not observe any influence of Rad21 on CTCF‐SA interactions, suggesting that SAs are stabilized by CTCF ZFs independently of other cohesin subunits. In agreement with this, Rad21‐independent colocalization of CTCF and SA have also been observed *in vivo*^23^. The direct recruitment of SA by CTCF ZFs as well as CTCF’s high stability on CBSs and ability to perform secondary RNA capture could therefore regulate SA1 and SA2’s different functions in transcription regulation and genome architecture^71,72,75^, possibly also by cohesin-independent interactions^23^.

## Conclusion

CTCF’s different binding modes on chromatin enable it to perform a variety of different functions inside the nucleus **(Figure 6)**. When binding unspecifically to DNA, CTCF displays a low residence time. CTCF can form oligomers, which can perform secondary capture of RNA, leading to more stable DNA binding. These oligomers could therefore act as an interaction hub for additional proteins like cohesin’s SA subunits^69^ or RNA Pol‐II^31^. Alternatively, monomeric CTCF can reach its target site by diffusion or rapid sampling of DNA, being stabilized on its CBS by more favorable ZF interactions. CBS-bound CTCF ZFs recruit SA by a Rad21-independent interaction, increasing SA’s residence time. CTCF can therefore influence cohesin-independent functions of SAs in the nucleus^23^. SAs may then recruit other cohesin subunits, facilitating TAD-formation. Alternatively, CTCF might block loop extrusion by its N‐terminal interaction with cohesin^21^ and subsequently increase loop-stability by the SA-ZF interaction. We conclude that the high-stability nucleic acid engagements of CTCF’s multiple ZFs enable its diverse roles in transcription regulation and TAD‐formation.

**Figure 6.**
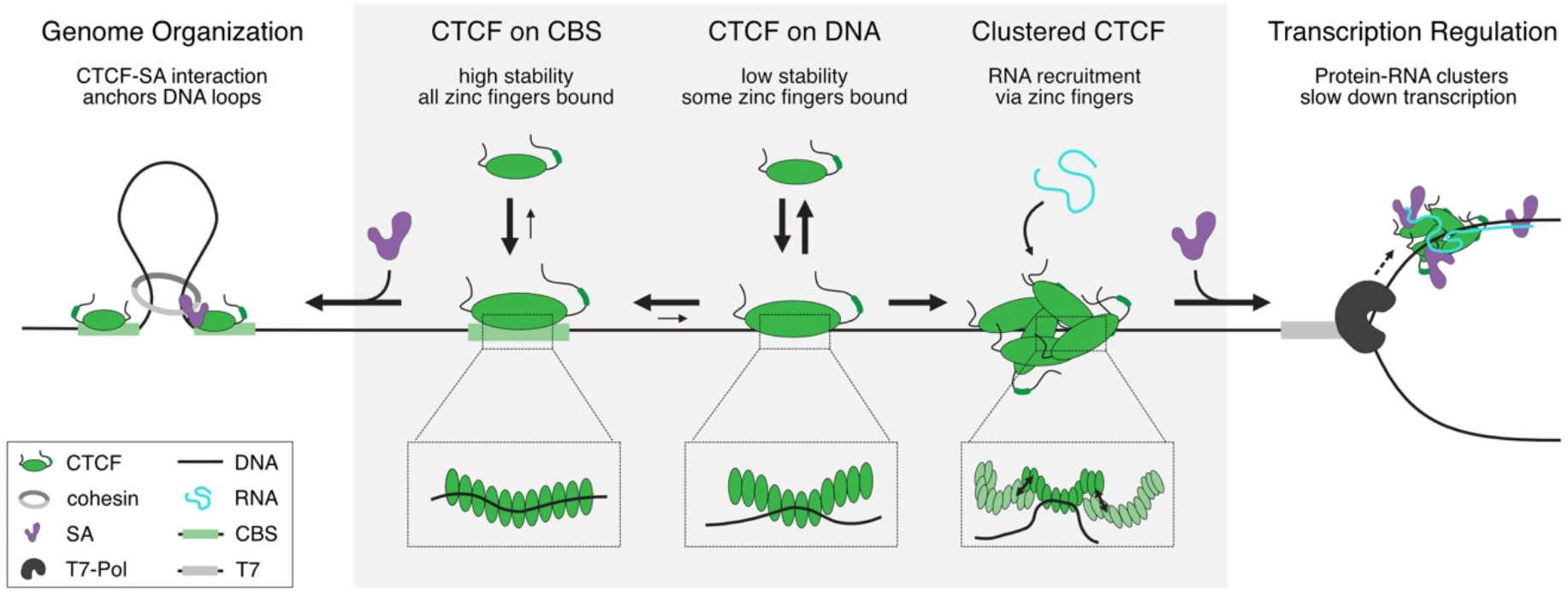
CTCF’s nucleic acid interactions regulate diverse processes in the nucleus.

## Materials and Methods

### CTCF expression, purification and labeling

Long CTCF constructs (CTCF WT, ΔN, ΔC, ΔNC, ΔRBR) were expressed in High Five (Hi5) insect cells. To this regard we cloned cassettes containing the respective CTCF variant with an N‐terminal 6xHis- and Halo‐tag and a C‐terminal Flag‐tag into a pLIB expression vector (kindly provided by Karl‐Peter Hopfner, LMU Munich). Expression vectors were transformed into DH10MultiBac cells to create a bacmid. Virus amplification was repeated two times in Sf9 cells, before transfecting 1 x 10^6^ High Five cells/ml with a 1:500 virus dilution. Protein expression continued for 3 days at 27 °C before harvesting the cells and freezing them in liquid N_2_. Cells were resuspended in CTCF resuspension buffer (25 mM Hepes pH 8.3, 100 mM NaCl, 5 % glycerol, 0.05 % tween, 100 μM ZnCl_2_, 10 mM imidazole, 1 mM TCEP, 1 mM PMSF, protease inhibitor tablet (Roche)) and sonicated for 90 sec. Short CTCF constructs (ZF4‐7, ZF9‐CT) were cloned into a pET28a vector containing an N‐terminal 6xHis‐tag and a C‐terminal Flag‐tag. The plasmid was transformed into *E*.*coli* Rosetta (DE3) and cells were grown to an OD of 0.6 before inducing expression with 1 mM IPTG for 16 h at 18 °C. Cells were harvested, resuspended in CTCF resuspension buffer and sonicated for 20 min. Lysates for long and short constructs were treated with 2000 units Pierce universal nuclease (Thermo Scientific) for 2 h at RT before increasing the salt concentration to 500 mM NaCl. Lysates were centrifuged for 1 h at 42 krpm and 4 °C. Supernatants were filtered through 0.22 μm PES filters and incubated with 3 ml Ni‐NTA beads (Macherey‐Nagel) for 45 minutes at 4 °C. The supernatant was applied to a gravity flow column and beads were washed with 100 ml of nickel CTCF wash buffer (25 mM Hepes pH 8.3, 1000 mM NaCl, 5 % Glycerol, 0.05 % Tween, 100 μM ZnCl_2_, 10 mM imidazole, 1 mM TCEP) before eluting with 10 ml nickel CTCF elution buffer (25 mM Hepes pH 8.3, 500 mM NaCl, 5 % Glycerol, 0.05 % Tween, 100 μM ZnCl_2_, 500 mM imidazole, 1 mM TCEP). CTCF constructs were dialyzed against CTCF 500 buffer (25 mM Hepes pH 8.3, 500 mM NaCl, 5 % Glycerol, 0.05 % Tween, 100 μM ZnCl2, 1 mM DTT) before further purification on a Cytiva HiTrap Heparin HP column with a 500 to 1500 mM NaCl gradient. CTCF fractions were pooled and concentrated to 500 μl. For labeling, chloroalkane linker containing fluorescent dyes were generated by click‐chemistry. 2.5 mM Halo‐DBCO (Iris‐Biotech) was incubated for 4 h at 37 °C with 7.5 mM AF568‐azide (Lumiprobe) or Atto 643 (ATTO‐TEC GmbH). Alternatively, Halo Tag Alexa Fluor 660 (Promega) was used. CTCF was treated with 10 x excess of dye for 15 minutes at RT followed by a final purification step on a Cytiva Superose 6 column in CTCF 500 buffer. CTCF fractions were pooled, aliquoted, flash frozen in liquid N2, stored at -80 °C and thawed directly before DNA‐curtains measurements.

### SA1/SA2 and Rad21 expression and purification

SA1 and SA2 were expressed in Hi5 insect cells. The cassettes containing the respective SA variant with an N‐terminal 10xHis‐tag and a C‐terminal ybbR‐tag were cloned into pLIB expression vectors. Expression and cell harvesting were performed as described for CTCF. Cells were resuspended in SA resuspension buffer (20 mM Hepes pH 7.5, 100 mM NaCl, 10 % Glycerol, 15 mM imidazole 1 mM TCEP, 1 mM PMSF, protease inhibitor tablet (Roche)) and sonicated for 90 sec. Lysates were treated with 2000 units Pierce universal nuclease (Thermo Scientific) for 45 minutes at RT before increasing the salt concentration to 300 mM NaCl. Lysates were centrifuged for 1 h at 42 krpm and 4 °C. Supernatants were filtered through 0.22 μm PES filters and incubated with 3 ml Ni‐NTA beads for 45 minutes at 4 °C. The supernatant was applied to a gravity flow column and beads were washed with 100 ml of nickel SA wash buffer (20 mM Hepes pH 7.5, 2000 mM NaCl, 10 % Glycerol, 40 mM imidazole 1 mM TCEP) before eluting with 10 ml nickel SA elution buffer (20 mM Hepes pH 7.5, 300 mM NaCl, 10 % Glycerol, 40 mM imidazole 1 mM TCEP). SA constructs were dialyzed for 1 h at 4 °C against SA 300 buffer (20 mM Hepes pH 7.5, 300 mM NaCl, 10 % Glycerol, 1 mM DTT) before further purification on a Cytiva HiTrap Heparin HP column with a 300 to 1000 mM NaCl gradient. SA fractions were pooled and concentrated to 500 μl. For ybbr‐tag labeling, Sfp (made in house, plasmid was kindly provided by the Gaub‐lab, LMU Munich) was mixed with SA in a molar ratio of 1:1 and a 1.25 excess of LD655‐CoA dye (Lumidyne) as well as 10 mM MgCl_2_. Reaction was performed for 16 h at 4 °C. Proteins were purified on a Cytiva Superose 6 column in SA 300 buffer. Protein fractions were pooled, frozen in liquid N_2_, stored at ‐80 °C and thawed directly before measurements. For SA‐Rad21 constructs, a Rad21 peptide consisting of amino acids 281-420 was fused to a N‐terminal 6xHis- and MBP‐tag in a pET28a vector and expressed in E. coli Rosetta (DE3) cells as described above for small CTCF constructs. Harvested cells were resuspended in SA resuspension buffer and sonicated for 20 min. Lysates were treated with 2000 units Pierce universal nuclease for 45 minutes at RT before increasing the salt concentration to 500 mM NaCl and centrifugation at 4 °C and 17 krpm for 30 min. Supernatants were filtered through 0.22 μm PES filters and incubated for 45 minutes at 4 °C with 3 ml Ni‐NTA beads. The supernatant was applied to a gravity flow column and beads were washed with 100 ml of nickel SA wash buffer before eluting with 10 ml nickel Rad21 elution buffer (40 mM Hep pH 7.5, 500 mM NaCl, 400 mM Imidazole, 5 % Glycerol, 1 mM TCEP). Proteins were dialyzed at 4 °C for 1 h against Rad21 500 buffer (40 mM Hep pH 7.5, 500 mM NaCl, 5 % Glycerol, 1 mM DTT) and concentrated to 500 μl. Rad21 and SA1 or SA2 (after Heparin) were then mixed in a molar ratio of 1:1 and filled up to 500 μl with SA 300 buffer. Heterodimers were labeled and purified like SA.

### Generation of λ-DNA constructs for DNA curtains

Wild‐type λ‐DNA was purchased from NEB. For generation of 2x T7‐4xCBSs λ‐DNA, a cassette containing a total of four 129 bp spaced CTCF binding sites (2x 5’‐TGCAGTTCCAAAACTGGCCAGCAGAGGGCACCAAA‐3’ and 2x 5’‐TGCAGTTCCAAAAGCGGCCAGCAGGGGGCGCCCAA‐3’) and a 2700 bp upstream T7 promoter site (5’‐TAATACGACTCACTATAGG‐3’) was cloned at two positions in opposite orientation into λ‐DNA using NgoMIV/XbaI and XhoI/NheI sites. For generation of 1xT7‐1xCBS λ‐DNA a single CTCF site (5’‐TGCAGTACCAACTTTAACCAGCAGAGGGCACCAAA‐3’) was cloned into the XhoI/NheI site and a single T7 promoter into the NgoMIV/XbaI site. The products were then packaged into phage particles using phage extract (MaxPlax, Epicentre) and amplified by lytic growth in LE392 cells (NEB). Following lytic growth, λ‐DNA was purified by PEG-precipitation and phenol-chloroform extraction before resuspension in TE 150 buffer (10 mM Tris pH 7.5, 1 mM EDTA, 150 mM NaCl). DNA ends were tagged with either biotin (5’‐aggtcgccgccc‐bio‐3’) or digoxigenin (5’‐gggcggcgacct‐dig‐3’) containing oligonucleotides (Metabion) by hybdridization and ligation to cos sites and purified on a HiPrep 16/60 Sephacryl S‐300 HR column in TE 150 buffer.

### Single-molecule DNA curtains experiments

DNA curtain experiments were carried out as described previously^77^ on a prism-type TIRF microscope (Nikon Eclipse Ti2), equipped with three illumination lasers (488 nm, 561 nm and 640 nm Coherent OBIS), an electron multiplying charged coupled camera (iXon Life, Andor) and a syringe-pump-driven microfluidics system supplying the sample chamber. Custom made flow cells were assembled from silica-fused slides grafted with chromium barriers produced via E‐beam lithography and cover slips with a double-sided tape.

Flow cells were incubated with a lipid mixture (Avanti) consisting of 10 mg/ml DOPC, 1 mg/ml DOPE‐PEG, 0.05 mg/ml DOPE‐biotin in lipid buffer (10 mM Tris pH 7.5, 200 mM NaCl, 20 mM MgCl_2_) in three 10 minutes incubation steps. After washout, flow cells were incubated with 2 μl 1 mg/ml anti‐digoxigenin (produced in house) in 700 μl lipid buffer followed by 5 μl streptavidin (Carl Roth) in 1 ml BSA buffer (40 mM Tris pH 7.5, 1 mg/ml BSA, 1 mM MgCl_2_). After this, 0.3 pM λ‐DNA in BSA buffer was added in four steps with 5 minutes incubation time. Single-molecule measurements were performed in CTCF 50 buffer (40 mM Tris pH 7.5, 1 mg/ml BSA, 1 mM MgCl_2_, 50 mM NaCl, 1 mM DTT) including an oxygen scavenger system containing glucose‐oxidase (Carl Roth), catalase (Sigma Aldrich) and 0.4 % glucose. Videos were recorded in NIS Elements (Nikon) and analyzed in Igor Pro 8 (Wavemetrics) using custom written code.

#### CTCF on DNA curtains

10 nM CTCF was incubated for 30 s on the DNA curtains to cover the DNA substrate, followed by CTCF enrichment on the CBS by a high salt buffer wash using CTCF 300 buffer (40 mM Tris pH 7.5, 1 mg/ml BSA, 1 mM MgCl_2_, 300 mM NaCl, 1 mM DTT) for 3 min. For this, CTCF 300 buffer was flushed in, and flow was then stopped for incubation time. For lifetime measurements, this incubation was increased to up to 60 minutes.

CTCF was imaged using a 561 nm laser at 50 mW for lifetime and diffusion videos and 140 mW for photobleaching analysis. For lifetime analysis, 100 ms illumination time and a frame delay of 1 s, 4 s, 10 s, 40 s or 60 s was used **(Figure S1A)**.

#### SA1 and SA2 on DNA curtains

100 nM SA1, SA2, SA1‐Rad21 or SA2‐Rad21 were incubated for 3 minutes on the flow cell in CTCF 50 buffer or CTCF 150 buffer and videos were recorded with 100 ms illumination times and 2 ms frame delay using a 640 nm laser at 140 mW.

#### SA and CTCF on DNA curtains

For sequential load experiments, first CTCF and then SA were loaded as described above and videos were recorded with 100 ms illumination times at 1s frame delay using a 561 nm laser at 50 mW and a 640 nm laser at 140 mW. For combined loading experiments, 100 nM SA was preincubated with 10 nM CTCF in CTCF 100 buffer (40 mM Tris pH 7.5, 1 mg/ml BSA, 1 mM MgCl_2_, 100 mM NaCl, 1 mM DTT) for 5 minutes at RT before 30 s incubation in CTCF 50 buffer on the DNA curtains and 3 minutes enrichment in CTCF 300 buffer. For lifetime measurements, incubation in CTCF 300 buffer was increased to up to 15 minutes.

#### Transcription experiments on DNA curtains

Transcription experiments were performed by incubation of 0.33 mM ATP, CTP, GTP, UTP (NEB), 0.03 mM Cy3-UTP (Jena Bioscience), and 1:300 T7 RNA polymerase mix (NEB) in transcription buffer (30 mM Tris pH 7.5, 2 mM spermidine, 25 mM MgCl_2_, 10 mM DTT) for 3 minutes at 40 °C, before loading for 2 minutes on DNA curtains. After wash-out, a 1 mM ATP, CTP, GTP, UTP (missing labeled UTP) solution in transcription buffer was loaded and a video was recorded at 100 ms illumination time and 1 s frame delay using a 561 nm laser with 25 mW laser power.

In case of CTCF transcription experiments, CTCF-Alexa Fluor 660 was loaded and enriched as described above before starting T7‐Pol transcription as described above. Transcription was recorded using 1 s frame delay and 25 mW of 561 nm laser to detect produced RNA and 10 s frame delay and 70 mW of 640 nm laser to visualize CTCF. For SA transcription experiments, SA‐LD655 was incubated with unlabeled CTCF and enriched on CBSs as described above. Then, the two-step transcription was started. Transcription was recorded using 1 s frame delay and 25 mW of 561 nm laser to detect RNA and 1 s frame delay and 140 mW of 640 nm laser to visualize SA.

#### RNA interaction on DNA curtains

For RNA recruitment experiments, 100 bp Cy3‐UTP labeled RNA was generated using a PCR‐product containing a T7‐promoter site and the HighScribe T7 high yield RNA synthesis kit (NEB) as well as Cy3‐UTP (Jena Bioscience). CTCF was incubated on DNA curtains with or without high salt wash and 25 ng/μl RNA in CTCF 50 buffer was added to the curtains and incubated for 3 minutes. CTCF was imaged using the 640 nm laser line at 70 mW and 10 s frame delay, RNA using the 561 nm laser line at 50 mW and 1 s frame delay. Photobleaching experiments were carried out with both laser lines at 140 mW and 100 ms frames with 2 ms frame delay. RNA recruitment to SA was performed accordingly. First, SA was incubated with DNA curtains as described above. Second, 25 ng/μl RNA in CTCF 50 buffer was added to the curtains and incubated for 3 minutes. SA1 was imaged using the 640 nm laser at 140 mW and 1 s frame delay.

For transcription bubble interaction experiments, transcription was performed on DNA curtains as described above. Afterwards T7‐Pol was washed off for 3 minutes in CTCF 1000 buffer (40 mM Tris pH 7.5, 1 mg/ml BSA, 1 mM MgCl_2_, 1000 mM NaCl, 1 mM DTT). SA or CTCF were then loaded as described above. Lifetime measurements were performed in CTCF 150 buffer for SAs and CTCF 300 buffer for CTCF.

### Data analysis for DNA curtains assay

#### Enrichment, survival and photobleaching data

DNA curtains data was analyzed in Igor Pro 8 using custom written code. Binding positions were determined in relation to positions of chromium anchors and barriers and divided into 40 bins. Enrichment on CTCF binding sites was calculated by determining the amount of CTCF or SA molecules in the bins containing CTCF binding sites in comparison to all other bins. Photobleaching data was analyzed according to a published method^78^.

SA survival data was calculated from the fraction of SA-molecules staying bound to DNA during a 3-minute wash.

#### Lifetimes

To determine lifetime data unbiased by photobleaching, the time of disappearance of fluorescent spots was recorded and turned into a survival curve using the Kaplan‐Meier method. As individual CTCF molecules could not be spatially separated on 4xCBSs constructs, a model was developed that describes the disappearance of the last molecule on a multi-binding-site array. The disappearance can be caused either by photobleaching or by dissociation from the binding site. The parameterλis hence given by

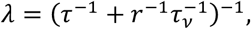

where *r* is the dimensionless ratio between off-times of the laser illumination and the observation times, and the parameter τ_*v*_ is the lifetime of the fluorescent dye until photobleaching at 100 % illumination. With this, the cumulative probability density function until the disappearance of the *i*‐th fluorophore from a fluorescent spot is

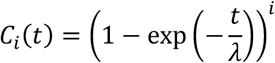

The fit model to the observed survival curve is then

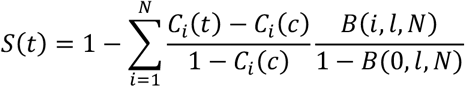

where *c* is the cut-off time after which the observation was started, *l* is the pre-determined labeling ratio and *N* is the number of binding sites (*N*= 4or *N* = 1in our case). The binomial distributions *B*(…) in the second term of the sum account for incomplete labeling. This fit model was verified using Monte Carlo simulations.

The experimental data was then fitted to this model with τ_*v*_ shared between all data sets, and *r* as common parameters for all measurements at the same frame rate.

The experimental data was then fitted to this model with the photobleaching rate τ_*v*_as a common parameter; the laser-off ratio *r* was set as a common parameter for all measurements at the same frame rate, the number of binding sites *N* and labeling ratio *l* as a common parameter for measurements with the same DNA, the true CTCF lifetime τ for measurements with the same construct and the shortest reliably measured lifetime *c* individually for each measurement. The lifetime of RNA and SA molecules were fitted to a single exponential model.

#### Movement analysis

For analysis of molecular movement, individual fluorescent spots were localized and tracked as described^79,80^. Molecules were split into two fractions (CBS-bound vs not CBS-bound for CTCF and bound at ≥ 50 % AT vs < 50 % AT for SA). Diffusion coefficients were determined according to a published method^81^.

Transcription speeds were determined from the onset of transcription until stalling/fall off or bleaching for each individual polymerase. Early stopping events were defined by transcription termination within the first 10 kbp; snapback events were defined as a sudden upward movement by at least 0.5 kbp. Polymerases that did not stop permanently were counted as continuously transcribing.

### Bulk Transcription Assay

DNA constructs for bulk transcription assays were generated by PCR from a pET28a vector or from λ‐DNA containing 4x CTCF binding sites followed by subsequent PCR purification. DNA constructs contained a T7‐promoter followed by either 100 bp DNA (no CTCF sites), 5093 bp DNA (4x CBSs upstream motif facing T7 first after 2672 bp) or 1039 bp DNA (4x CBSs opposite orientation first after 196 bp). Reactions using equimolar amount of DNA were carried out in presence or absence of 50 nM CTCF using a HiScribe T7 High Yield RNA Synthesis Kit (NEB), adding 100 μM Cy3‐UTP. Following DNAse treatment, products were purified using an RNA purification kit (NEB), absorption at 260 nm was determined and length of the RNA product was analyzed on a 1 % agarose gel.

### Mass Photometry

Mass photometry measurements were performed on a Refeyn OneMP mass photometer with an image size of 10.8 μm x 2.9 μm. Cover slips (Roth) were cleaned by sonication in isopropanol and dried with nitrogen. Silicon culture well gaskets (Merck) were placed on the slide and 19 μl buffer was pipetted into the well. After focusing the laser, 1 μl protein was added to the buffer droplet to a final concentration of 50 nM and mixed by pipetting. Measurement was performed for 1 minute using Acquire software (Refeyn). For data analysis, Discover software (Refeyn) was used to convert measured contrasts into molecular masses (calibration was done using protein standards with known molecular mass). A histogram with 100 bins was calculated from all measured masses and Gaussian fits were applied to individual protein peaks using Igor Pro 8 (Wavemetrics).

### Statistical Analysis

Lifetime data and SA diffusion data were analyzed using a two-tailed z-test. Counted single-molecule events for passing/blocking/binding SA or CTCF molecules as well as processive/impaired transcription events were analyzed using fisher’s exact test. All other data was analyzed using a two-tailed t-test. A table containing number of molecules and p-values for all relevant experiments can be found in the supplement.

## Supporting information

Supplementary information (figures/statistical tests/protein sequences)

## Data availability

The data that supports the findings of our study is available upon reasonable request.

## Author Contributions

J.H. conducted the experiments. S.Z. conducted mass photometry experiments. N-L.T. and J.H. established the single-molecule transcription assay. J.H. and J.S. analyzed the data. N.-L.T. analyzed CTCF diffusion data. All authors wrote the paper. J.S. designed the study, provided funding and supervised the project.

## Acknowledgments

We thank Marvin Freitag, Jeanny Probst, Joelle Deplazes-Lauber and Sigrun Jaklin for preparation of λ-DNA constructs. Thanks also to Bingzhi Wang for preparing a Python script for single-molecule photobleaching analysis, and Daniel Bollschweiler for providing access to mass photometry. We thank Prof. Christof Gebhardt for providing the CTCF WT construct. Thanks also to all members of the Stigler lab for helpful discussions. This work was supported by the LMU Center for Nanoscience CeNS and an ERC starting grant (no. 758124).

