## Supplementary information (figures/statistical tests/protein sequences) for "Single-molecule imaging reveals a direct role of CTCF’s zinc fingers in SA interaction and cluster-dependent RNA recruitment"

This file contains:

**Supplementary figures**

**Significance tests**

**Protein sequences**

### Supplementary figures

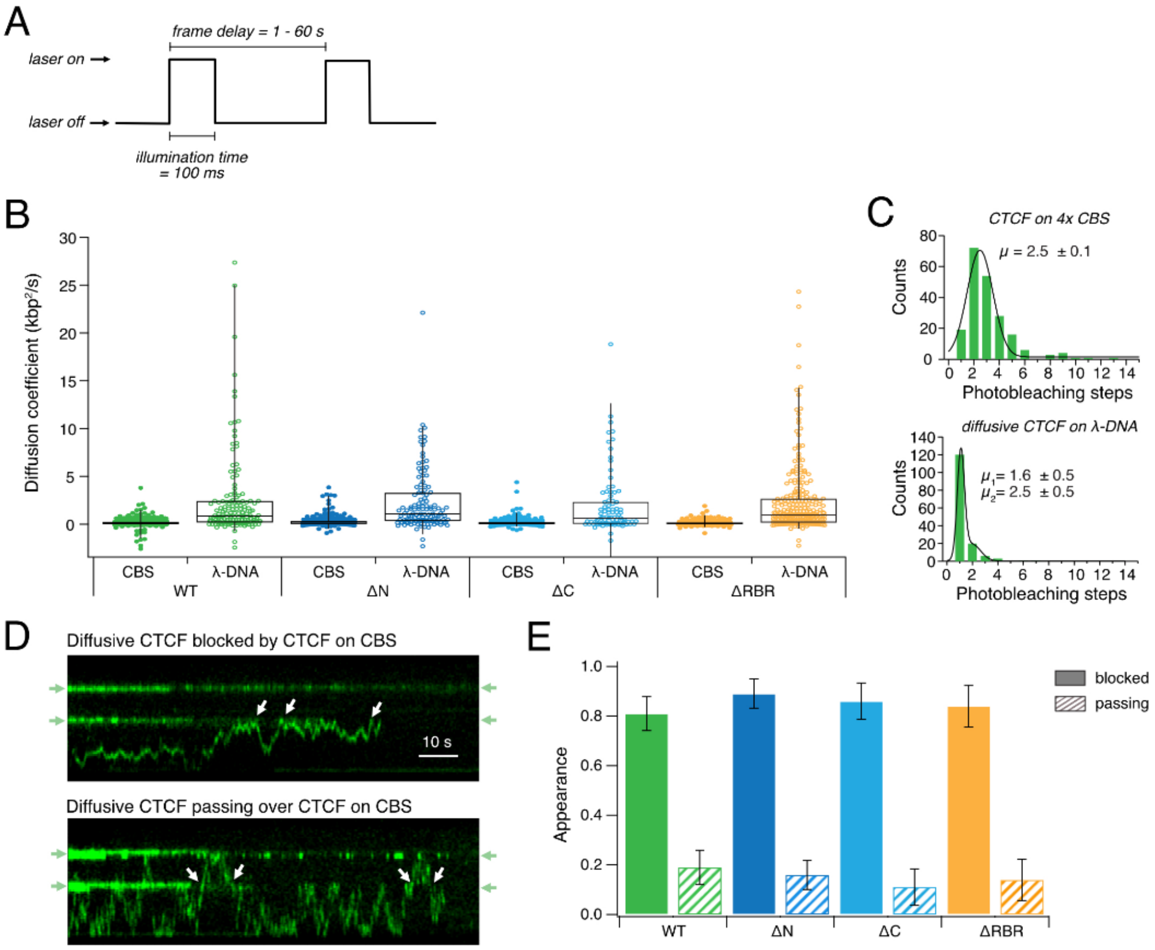


**Figure S1.** CTCF diffuses on non-CBS sites. **(A)** Scheme illustrating laser illumination during lifetime measurements. **(B)** Diffusion coefficients of CTCF WT and variants on 4x CBSs or λ-DNA. For all variants, D is significantly higher on λ-DNA than CBSs. No significant difference in diffusive behavior between CTCF variants. **(C)** Photobleaching steps of non-diffusive (top) and diffusive (bottom) CTCF. **(D)** Representative kymograms showing diffusion behavior of WT CTCF. Top: White arrows indicate events where diffusive CTCF is blocked by CBS-bound CTCF. No recruitment of diffusive CTCF to the binding sites occurs. Bottom: White arrows indicate events where diffusive CTCF passes CBS-bound CTCF. **(E)** Quantification of blocking and passing events. No significant differences were observed between WT and CTCF variants.

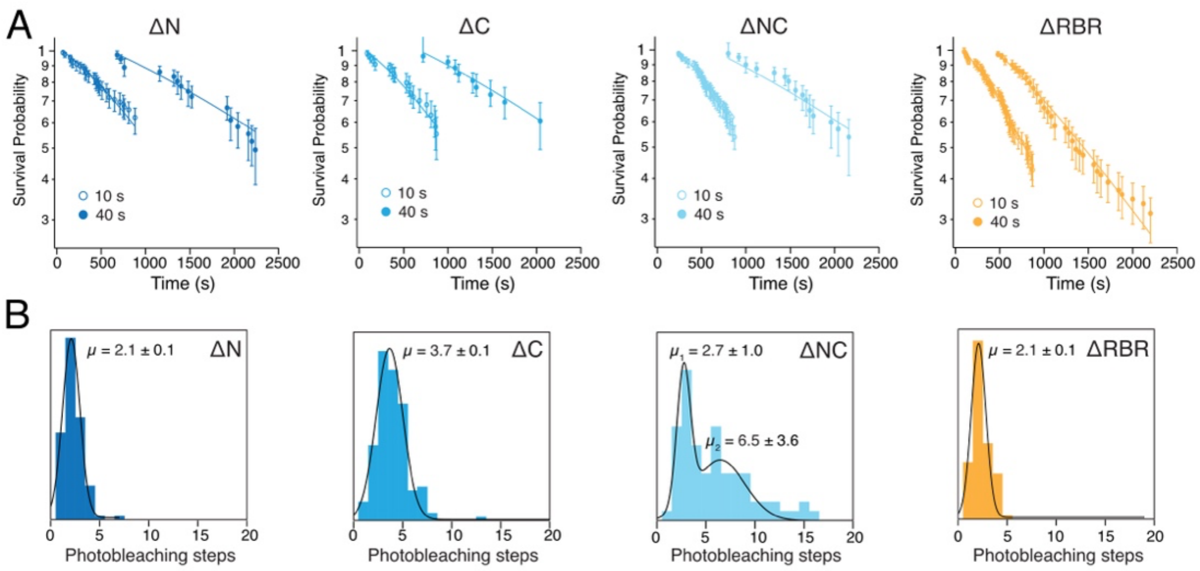


**Figure S2.** Lifetimes of CTCF variants. **(A)** Lifetimes of CTCF variants at 10 and 40 s frame rates. **(B)** Photobleaching steps of CTCF variants binding to 4x CBSs. Black line: Multi-Gaussian fit.

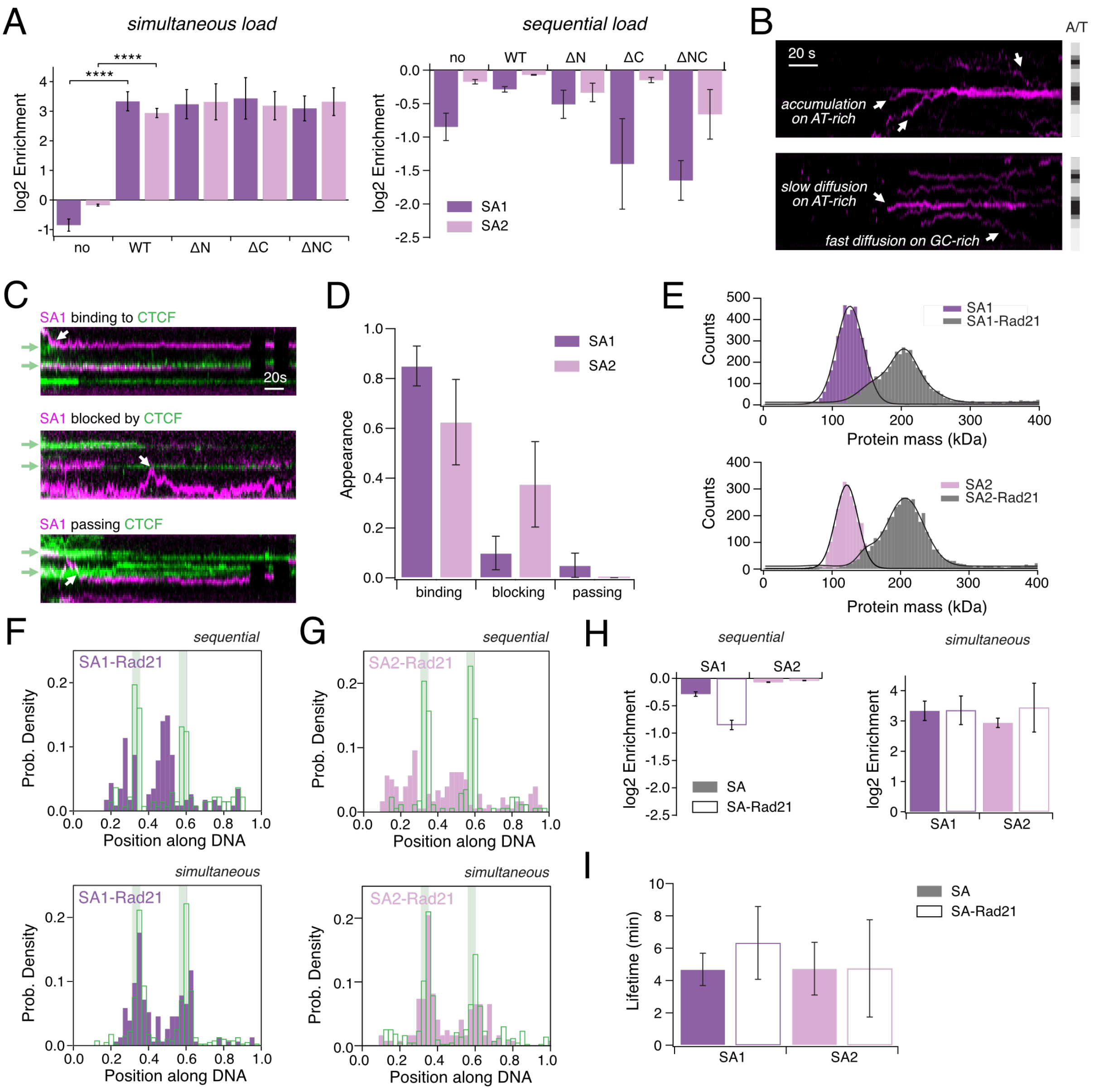


**Figure S3.** SA diffusion behavior and influence of Rad21 on CTCF-SA interaction. **(A)** Enrichment of SA1 (purple) and SA2 (pink) on CBSs. During simultaneous load (left), both SAs are enriched significantly more on CBSs when preincubated with CTCF WT compared to SAs loaded alone. SA is enriched similarly in presence of all CTCF variants. When CTCF was bound first to DNA, followed by SA (sequential load, right), no significant SA enrichment at CBSs was observed with any CTCF variant. **(B)** Representative kymograms of SAs binding static to AT-rich and diffusing randomly on GC-rich regions. **(C)** Representative kymograms of binding (top), blocking (middle) and passing (bottom) events observed for diffusive SAs (magenta) after sequential load on CTCF (green). **(D)** Fraction of binding, blocking and passing events observed for SA1 and SA2 after sequential load. No significant difference was found between SA1 and SA2. **(E)** Mass photometry data of 148 kDa SA1 (top, purple) and 145 kDa SA2 (bottom, pink) in absence and presence (gray) of 59 kDa Rad21-MBP. **(F)** Histogram of SA1-Rad21 and CTCF binding positions for sequential load (top, CTCF, N = 137; SA1-Rad21, N = 114) and simultaneous load (bottom, CTCF, N = 397; SA1-Rad21, N = 153) experiments. **(G)** Same as (E) but using SA2-Rad21 (sequential, CTCF, N = 292; SA2-Rad21, N = 257; simultaneous, CTCF, N = 433; SA2-Rad21, N = 122). **(H)** Enrichment of SAs on CBSs in absence (solid) or presence (transparent) of Rad21 for sequential (left) and simultaneous load (right). Enrichments are independent from Rad21. **(I)** Lifetimes of SAs on CTCF in absence (solid) and presence (transparent) of Rad21. Lifetimes are independent of Rad21.

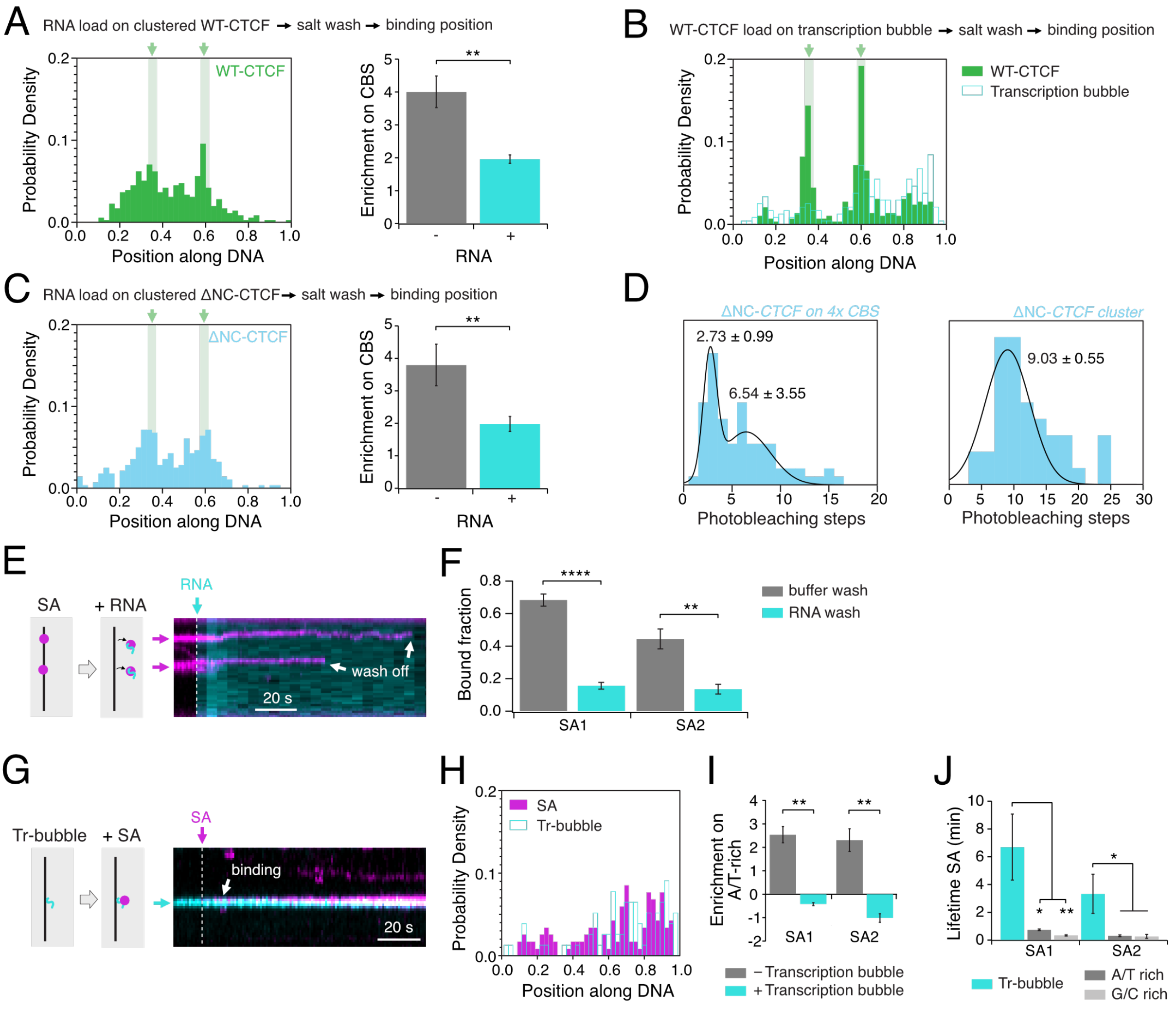


**Figure S4**. SA has a higher affinity for RNA than for DNA. **(A)** Left: Histogram of CTCF WT-RNA cluster (N = 355) binding positions after salt enrichment (see Figure 5D). Right: CTCF WT-RNA clusters are significantly less enriched on CBSs than monomeric CTCF WT. **(B)** Histogram of CTCF WT binding positions after transcription and salt enrichment (see Figure 5G). CTCF preferentially binds to 4x CBSs but also colocalizes with transcription bubbles (CTCF, N = 292; RNA, N = 237). **(C)** Left: Histogram of CTCF ΔNC-RNA cluster (N = 280) binding positions after salt enrichment. Right: ΔNC-RNA clusters are significantly less enriched on CBSs than monomeric ΔNC. **(D)** Histogram of photobleaching steps for ΔNC on 4x CBSs (same as Figure S2B-3) and in RNA-clusters. **(E)** Scheme and representative kymogram of SAs being washed off from λ-DNA by the addition of RNA to the DNA-curtain. **(F)** More SA1 and SA2 is washed off in the presence of RNA than in the presence of buffer. **(G)** Scheme and representative kymogram of SAs colocalizing with transcription bubbles. **(H)** Histogram of SA1 binding positions after transcription (SA1, N = 43; RNA, N = 155). **(I)** SA1 and SA2 are significantly less enriched on AT-rich regions in presence of transcription bubbles on the DNA. **(J)** SA1 and SA2 have a significantly higher lifetime on transcription bubbles than on AT-rich or GC-rich DNA regions.

**Significance tests**

Two tailed t-test: (t)

Two tailed z-test: (z)

Fisher's exact test: (f)

| | p-values | N |
|---|---|---|
| **Figure 1** | | |
| (F) 4x CBSs/1x CBS | 0.016 (t) | 427/477 |
| (H) 4x CBSs/1x CBS | 0.89 (z) | 701/201 |
| (H) 4x CBSs/λ-DNA | $< 10^{-6}$ (z) | 701/271 |
| (H) 1x CBS/λ-DNA | $< 10^{-6}$ (z) | 201/271 |
| **Figure S1** | | |
| (B) λ-DNA /CBS WT | $2.6 * 10^{-6}$ (t) | 148/261 |
| (B) λ-DNA /CBS ΔN | $< 10^{-6}$ (t) | 125/235 |
| (B) λ-DNA /CBS ΔC | $1.4 * 10^{-6}$ (t) | 89/288 |
| (B) λ-DNA / CBS ΔRBR | $< 10^{-6}$ (t) | 266/189 |
| (B) WT/ΔN λ-DNA | 0.37 (t) | 148/125 |
| (B) WT/ΔC λ-DNA | 0.19 (t) | 148/89 |
| (B) WT/ΔRBR λ-DNA | 0.24 (t) | 148/266 |
| (E) WT/ΔN | 0.48 (f) | 32/28 |
| (E) WT/ΔC | 0.72 (f) | 32/22 |
| (E) WT/ΔRBR | 1 (f) | 32/19 |
| **Figure 2** | | |
| (B) WT/ΔN | 0.22 (t) | 427/390 |
| (B) WT/ΔC | 0.63 (t) | 427/666 |
| (B) WT/ΔNC | 0.49 (t) | 427/396 |
| (B) WT/ΔRBR | 0.26 (t) | 427/651 |
| (B) WT/ZF9-CT | 0.0012 (t) | 427/343 |
| (B) WT/ZF4-7 | 0.0020 (t) | 427/88 |
| (D) WT/ΔN | 0.97 (z) | 701/125 |
| (D) WT/ΔC | 0.93 (z) | 701/80 |
| (D) WT/ΔNC | 0.89 (z) | 701/161 |
| (D) WT/ΔRBR | $3.4 * 10^{-6}$ (z) | 701/217 |
| (D) WT/ZF9-CT | $< 10^{-6}$ (z) | 701/155 |
| (D) WT/ZF4-7 | $< 10^{-6}$ (z) | 701/88 |
| **Figure 3** | | |
| (G) 50 mM NaCl SA1/SA2 GC | 0.012 (z) | 186/48 |
| (G) 50 mM NaCl SA1/SA2 AT | 0.37 (z) | 54/62 |
| (G) 50 mM NaCl GC/AT SA1 | $< 10^{-6}$ (z) | 54/186 |
| (G) 50 mM NaCl GC/AT SA2 | 0.0019 (z) | 62/48 |
| (G) 150 mM NaCl SA1/SA2 GC | 0.63 (z) | 8/20 |
| (G) 150 mM NaCl SA1/SA2 AT | $1.2 * 10^{-6}$ (z) | 65/62 |
| (G) 150 mM NaCl GC/AT SA1 | $< 10^{-6}$ (z) | 8/65 |
| (G) 150 mM NaCl GC/AT SA2 | 0.77 (z) | 20/62 |

| | | |
|---|---|---|
| (H) SA1/SA2 GC | 0.0058 (z) | 537/510 |
| (H) SA1/SA2 AT | 0.92 (z) | 231/169 |
| (H) GC/AT SA1 | $2.1 * 10^{-5}$ (z) | 537/231 |
| (H) GC/AT SA2 | $< 10^{-6}$ (z) | 510/169 |
| (I) WT/DNA SA1 | $7.7 * 10^{-5}$ (z) | 65/8 |
| (I) WT/DNA SA2 | 0.0044 (z) | 62/20 |
| (I) WT/ ΔN SA1 | 0.41 (z) | 65/28 |
| (I) WT/ΔC SA1 | 0.62 (z) | 65/20 |
| (I) WT/ΔNC SA1 | 0.76 (z) | 65/9 |
| (I) WT/ΔN SA2 | 0.83 (z) | 62/28 |
| (I) WT/ΔC SA2 | 0.66 (z) | 62/17 |
| (I) WT/ΔNC SA2 | 0.85 (z) | 62/10 |
| **Figure S3** | | |
| (A)WT/DNA SA1 simultaneous | $2.5 * 10^{-4}$ (t) | 332/1321 |
| (A)WT/DNA SA2 simultaneous | $3.3 * 10^{-6}$ (t) | 224/875 |
| (A)WT/ΔN SA1 simultaneous | 0.74 (t) | 332/197 |
| (A)WT/ΔN SA2 simultaneous | 0.31 (t) | 224/153 |
| (A)WT/ΔC SA1 simultaneous | 0.79 (t) | 332/251 |
| (A)WT/ΔC SA2 simultaneous | 0.41 (t) | 224/205 |
| (A)WT/ΔNC SA1 simultaneous | 0.39 (t) | 332/116 |
| (A)WT/ΔNC SA2 simultaneous | 0.22 (t) | 224/40 |
| (A)WT/DNA SA1 sequential | 0.22 (t) | 380/1321 |
| (A)WT/DNA SA2 sequential | 0.76 (t) | 80/875 |
| (A)WT/ΔN SA1 sequential | 0.73 (t) | 380/114 |
| (A)WT/ΔN SA2 sequential | 0.85 (t) | 80/88 |
| (A)WT/ΔC SA1 sequential | 0.55 (t) | 380/265 |
| (A)WT/ΔC SA2 sequential | 0.97 (t) | 80/242 |
| (A)WT/ΔNC SA1 sequential | 0.18 (t) | 380/207 |
| (A)WT/ΔNC SA2 sequential | 0.63 (t) | 80/95 |
| (D)SA1/SA2 binding | 0.31 (f) | 20/8 |
| (D)SA1/SA2 blocking | 0.12 (f) | 20/8 |
| (D)SA1/SA2 passing | 1.0 (f) | 20/8 |
| (G)SA1/SA1Rad21 on WT simultaneous | 0.95 (t) | 332/137 |
| (G)SA2/SA2Rad21 on WT simultaneous | 0.31 (t) | 224/277 |
| (G)SA1/SA1Rad21 on WT sequential | 0.15 (t) | 380/153 |
| (G)SA2/SA2Rad21 on WT sequential | 0.90 (t) | 80/122 |
| (H)SA1/SA1Rad21 on WT | 0.51 (z) | 65/16 |
| (H)SA2/SA2Rad21 on WT | 1.0 (z) | 62/9 |
| **Figure 4** | | |
| (A)CTCF/no | 0.78 (t) | 3/3 |
| (B)CTCF/no | 0.18 (t) | 3/3 |
| (F)T7/WT | 0.034 (t) | 111/88 |
| (F)T7/ΔN | 0.22 (t) | 111/84 |
| (F)T7/ΔC | 0.89 (t) | 111/76 |
| (F)T7/WT-SA1 | 0.63 (t) | 111/9 |

| | | |
|---|---|---|
| (F)T7/WT-SA2 | 0.85 (t) | 111/9 |
| (H)T7single/T7multiple | $< 10^{-6}$ (f) | 49/118 |
| (H)T7single/WTsingle | $< 10^{-6}$ (f) | 49/40 |
| (H)T7single/WTmultiple | $< 10^{-6}$ (f) | 49/147 |
| (H)T7single/ΔNsingle | $4.1 * 10^{-6}$ (f) | 49/33 |
| (H)T7single/ΔNmultiple | $< 10^{-6}$ (f) | 49/134 |
| (H)T7single/ΔCsingle | $< 10^{-6}$ (f) | 49/34 |
| (H)T7single/ΔCmultiple | $< 10^{-6}$ (f) | 49/177 |
| (H)T7multiple/WTmultiple | 0.0056 (f) | 118/147 |
| (H)T7multiple/ΔNmultiple | 0.0031 (f) | 118/134 |
| (H)T7multiple/ΔCmultiple | 0.042 (f) | 118/177 |
| **Figure 5** | | |
| (H)RNA/noRNA enrichment 4x CBSs | 0.060 (t) | 292/427 |
| (I) RNA/4x CBSs | $< 10^{-6}$ (z) | 118/201 |
| (I) RNA/λ-DNA | 0.11 (z) | 118/274 |
| **Figure S4** | | |
| (A)RNAcluster/monomers WT | 0.0021 (t) | 355/427 |
| (C)RNAcluster/monomers ΔNC | 0.0022 (t) | 280/396 |
| (F)RNA/noRNA wash off SA1 | $< 10^{-6}$ (t) | 560/479 |
| (F)RNA/noRNA wash off SA2 | 0.00267 (t) | 148/198 |
| (I)RNA/noRNA AT-enrichment SA1 | 0.0029 (t) | 1321/43 |
| (I)RNA/noRNA AT-enrichment SA2 | 0.0088 (t) | 875/117 |
| (J)RNA/AT | 0.012 (z) | 420/260 |
| (J)RNA/AT | 0.032 (z) | 295/119 |
| (J)RNA/GC | 0.0075 (z) | 420/93 |
| (J)RNA/GC | 0.030 (z) | 295/146 |

81

**Protein Sequences**

<u>CTCF WT</u>

6xHis – Halo – TEV site – CTCF WT – Flag

MGSSHHHHHHSSGTSLYKKAGLMAEIGTGFPFDPHYVEVLGERMHYVDVGPRDGTPVLFLHG
NPTSSYVWRNIIPHVAPTHRCIAPDLIGMGKSDKPDLGYFFDDHVRFMDAFIEALGLEEVVLVIHD
WGSALGFHWAKRNPERVKGIAFMEFIRPIPTWDEWPEFARETFQAFRTTDVGRKLIIDQNVFIEG
TLPMGVVRPLTEVEMDHYREPFLNPVDREPLWRFPNELPIAGEPANIVALVEEYMDWLHQSPV
PKLLFWGTPGVLIPPAEAARLAKSLPNCKAVDIGPGLNLLQEDNPDLIGSEIARWLSTLEISGEPT
TEDLYFQSDNTTLYTKVVMEGDAVEAIVEESETFIKGKERKTYQRRREGGQEEDACHLPQNQTD
GGEVVQDVNSSVQMVMMEQLDPTLLQMKTEVMEGTVAPEAEAAVDDTQIITLQVVNMEEQPINI
GELQLVQVPVPVTVPVATTSVEELQGAYENEVSKEGLAESEPMICHTLPLPEGFQVVKVGANGE
VETLEQGELPPQEDPSWQKDPDYQPPAKKTKKTKKSKLRYTEEGKDVDVSVYDFEEEQQEGLL
SEVNAEKVVGNMKPPKPTKIKKKGVKKTFQCELCSYTCPRRSNLDRHMKSHTDERPHKCHLCG
RAFRTVTLLRNHLNTHTGTRPHKCPDCDMAFVTSGELVRHRRYKHTHEKPFKCSMCDYASVEV
SKLKRHIRSHTGERPFQCSLCSYASRDTYKLKRHMRTHSGEKPYECYICHARFTQSGTMKMHIL
QKHTENVAKFHCPHCDTVIARKSDLGVHLRKQHSYIEQGKKCRYCDAVFHERYALIQHQKSHKN
EKRFKCDQCDYACRQERHMIMHKRTHTGEKPYACSHCDKTFRQKQLLDMHFKRYHDPNFVPA
AFVCSKCGKTFTRRNTMARHADNCAGPDGVEGENGGETKKSKRGRKRKMRSKKEDSSDSEN
AEPDLDDNEDEEEPAVEIEPEPEPQPVTPAPPPAKKRRGRPPGRTNQPKQNQPTAIIQVEDQNT
GAIENIIVEVKKEPDAEPAEGEEEEAQPAATDAPNGDLTPEMILSMMDRDYKDDDDK

<u>CTCF ΔN</u>

6xHis – Halo – TEV site – CTCF (Δ1-265) – Flag

MGSSHHHHHHSSGMAEIGTGFPFDPHYVEVLGERMHYVDVGPRDGTPVLFLHGNPTSSYVWR
NIIPHVAPTHRCIAPDLIGMGKSDKPDLGYFFDDHVRFMDAFIEALGLEEVVLVIHDWGSALGFH
WAKRNPERVKGIAFMEFIRPIPTWDEWPEFARETFQAFRTTDVGRKLIIDQNVFIEGTLPMGVVR
PLTEVEMDHYREPFLNPVDREPLWRFPNELPIAGEPANIVALVEEYMDWLHQSPVPKLLFWGTP
GVLIPPAEAARLAKSLPNCKAVDIGPGLNLLQEDNPDLIGSEIARWLSTLEISGEPTTEDLYFQSG
SFQCELCSYTCPRRSNLDRHMKSHTDERPHKCHLCGRAFRTVTLLRNHLNTHTGTRPHKCPDC
DMAFVTSGELVRHRRYKHTHEKPFKCSMCDYASVEVSKLKRHIRSHTGERPFQCSLCSYASRD
TYKLKRHMRTHSGEKPYECYICHARFTQSGTMKMHILQKHTENVAKFHCPHCDTVIARKSDLGV
HLRKQHSYIEQGKKCRYCDAVFHERYALIQHQKSHKNEKRFKCDQCDYACRQERHMIMHKRTH
TGEKPYACSHCDKTFRQKQLLDMHFKRYHDPNFVPAAFVCSKCGKTFTRRNTMARHADNCAG
PDGVEGENGGETKKSKRGRKRKMRSKKEDSSDSENAEPDLDDNEDEEEPAVEIEPEPEPQPV
TPAPPPAKKRRGRPPGRTNQPKQNQPTAIIQVEDQNTGAIENIIVEVKKEPDAEPAEGEEEEAQP
AATDAPNGDLTPEMILSMMDRDYKDDDDK

<u>CTCF ΔC</u>

6xHis – Halo – TEV site – CTCF (Δ580-727) – Flag

MGSSHHHHHHSSGMAEIGTGFPFDPHYVEVLGERMHYVDVGPRDGTPVLFLHGNPTSSYVWR
NIIPHVAPTHRCIAPDLIGMGKSDKPDLGYFFDDHVRFMDAFIEALGLEEVVLVIHDWGSALGFH
WAKRNPERVKGIAFMEFIRPIPTWDEWPEFARETFQAFRTTDVGRKLIIDQNVFIEGTLPMGVVR
PLTEVEMDHYREPFLNPVDREPLWRFPNELPIAGEPANIVALVEEYMDWLHQSPVPKLLFWGTP
GVLIPPAEAARLAKSLPNCKAVDIGPGLNLLQEDNPDLIGSEIARWLSTLEISGEPTTEDLYFQSD
NTTLYTKVVMEGDAVEAIVEESETFIKGKERKTYQRRREGGQEEDACHLPQNQTDGGEVVQDV
NSSVQMVMMEQLDPTLLQMKTEVMEGTVAPEAEAAVDDTQIITLQVVNMEEQPINIGELQLVQV
PVPVTVPVATTSVEELQGAYENEVSKEGLAESEPMICHTLPLPEGFQVVKVGANGEVETLEQGE

LPPQEDPSWQKDPDYQPPAKKTKKTKKSKLRYTEEGKDVDVSVYDFEEEQQEGLLSEVNAEKVVGNMKPPKPTKIKKKGVKKTFQCELCSYTCPRRSNLDRHMKSHTDERPHKCHLCGRAFRTVTLLRNHLNTHTGTRPHKCPDCDMAFVTSGELVRHRRYKHTHEKPFKCSMCDYASVEVSKLKRHIRSHTGERPFQCSLCSYASRDTYKLKRHMRTHSGEKPYECYICHARFTQSGTMKMHILQKHTENVAKFHCPHCDTVIARKSDLGVHLRKQHSYIEQGKKCRYCDAVFHERYALIQHQKSHKNEKRFKCDQCDYACRQERHMIMHKRTHTGEKPYACSHCDKTFRQKQLLDMHFKRYHDPNFVPAAFVCSKCGKTFTRRNTMARHADNCAGDYKDDDDK

<u>CTCF ΔNC</u>

6xHis – Halo – TEV site – CTCF (Δ1-265; Δ580-727) – Flag

MGSSHHHHHHSSGMAEIGTGFPFDPHYVEVLGERMHYVDVGPRDGTPVLFLHGNPTSSYVWRNIIPHVAPTHRCIAPDLIGMGKSDKPDLGYFFDDHVRFMDAFIEALGLEEVVLVIHDWGSALGFHWAKRNPERVKGIAFMEFIRPIPTWDEWPEFARETFQAFRTTDVGRKLIIDQNVFIEGTLPMGVVRPLTEVEMDHYREPFLNPVDREPLWRFPNELPIAGEPANIVALVEEYMDWLHQSPVPKLLFWGTPGVLIPPAEAARLAKSLPNCKAVDIGPGLNLLQEDNPDLIGSEIARWLSTLEISGEPTTEDLYFQSGSFQCELCSYTCPRRSNLDRHMKSHTDERPHKCHLCGRAFRTVTLLRNHLNTHTGTRPHKCPDCDMAFVTSGELVRHRRYKHTHEKPFKCSMCDYASVEVSKLKRHIRSHTGERPFQCSLCSYASRDTYKLKRHMRTHSGEKPYECYICHARFTQSGTMKMHILQKHTENVAKFHCPHCDTVIARKSDLGVHLRKQHSYIEQGKKCRYCDAVFHERYALIQHQKSHKNEKRFKCDQCDYACRQERHMIMHKRTHTGEKPYACSHCDKTFRQKQLLDMHFKRYHDPNFVPAAFVCSKCGKTFTRRNTMARHADNCAGDYKDDDDK

<u>CTCF ΔRBR</u>

6xHis – Halo – TEV site – CTCF (Δ264-291; Δ521-614) – Flag

MGSSHHHHHHSSGTSLYKKAGLMAEIGTGFPFDPHYVEVLGERMHYVDVGPRDGTPVLFLHGNPTSSYVWRNIIPHVAPTHRCIAPDLIGMGKSDKPDLGYFFDDHVRFMDAFIEALGLEEVVLVIHDWGSALGFHWAKRNPERVKGIAFMEFIRPIPTWDEWPEFARETFQAFRTTDVGRKLIIDQNVFIEGTLPMGVVRPLTEVEMDHYREPFLNPVDREPLWRFPNELPIAGEPANIVALVEEYMDWLHQSPVPKLLFWGTPGVLIPPAEAARLAKSLPNCKAVDIGPGLNLLQEDNPDLIGSEIARWLSTLEISGEPTTEDLYFQSDNTTLYTKVVMEGDAVEAIVEESETFIKGKERKTYQRRREGGQEEDACHLPQNQTDGGEVVQDVNSSVQMVMMEQLDPTLLQMKTEVMEGTVAPEAEAAVDDTQIITLQVVNMEEQPINIGELQLVQVPVPVTVPVATTSVEELQGAYENEVSKEGLAESEPMICHTLPLPEGFQVVKVGANGEVETLEQGELPPQEDPSWQKDPDYQPPAKKTKKTKKSKLRYTEEGKDVDVSVYDFEEEQQEGLLSEVNAEKVVGNMKPPKPTKIKKKGVKRPHKCHLCGRAFRTVTLLRNHLNTHTGTRPHKCPDCDMAFVTSGELVRHRRYKHTHEKPFKCSMCDYASVEVSKLKRHIRSHTGERPFQCSLCSYASRDTYKLKRHMRTHSGEKPYECYICHARFTQSGTMKMHILQKHTENVAKFHCPHCDTVIARKSDLGVHLRKQHSYIEQGKKCRYCDAVFHERYALIQHQKSHKNEKRFKCDQCDYACRQERHMIMHKRTHTGEAEPDLDDNEDEEEPAVEIEPEPEPQPVTPAPPPAKKRRGRPPGRTNQPKQNQPTAIIQVEDQNTGAIENIIVEVKKEPDAEPAEGEEEEAQPAATDAPNGDLTPEMILSMMDRDYKDDDDK

<u>ZF4-7</u>

6xHis – CTCF (Δ1-350; Δ461-727) – Flag

MGSSHHHHHHSSGFKCSMCDYASVEVSKLKRHIRSHTGERPFQCSLCSYASRDTYKLKRHMRTHSGEKPYECYICHARFTQSGTMKMHILQKHTENVAKFHCPHCDTVIARKSDLGVHLRKQHDYKDDDDK

<u>ZF9-CT</u>

6xHis – CTCF (Δ1-489) – Flag

MGSSHHHHHHSSGLVPRGSHMKNEKRFKCDQCDYACRQERHMIMHKRTHTGEKPYACSHCDKTFRQKQLLDMHFKRYHDPNFVPAAFVCSKCGKTFTRRNTMARHADNCAGPDGVEGENGGETKKSKRGRKRKMRSKKEDSSDSENAEPDLDDNEDEEEPAVEIEPEPEPQPVTPAPPPAKKRRGRPPGRTNQPKQNQPTAIIQVEDQNTGAIENIIVEVKKEPDAEPAEGEEEEAQPAATDAPNGDLTPEMILSMMDRDYKDDDDK

<u>SA1</u>

10xHis – SA1 WT – S6

MHHHHHHHHHHSGGSMITSELPVLQDSTNETTAHSDAGSELEETEVKGKRKRGRPGRPPSTNKKPRKSPGEKSRIEAGIRGAGRGRANGHPQQNGEGEPVTLFEVVKLGKSAMQSVVDDWIESYKQDRDIALLDLINFFIQCSGCRGTVRIEMFRNMQNAEIIRKMTEEFDEDSGDYPLTMPGPQWKKFRSNFCEFIGVLIRQCQYSIIYDEYMMDTVISLLTGLSDSQVRAFRHTSTLAAMKLMTALVNVALNLSIHQDNTQRQYEAERNKMIGKRANERLELLLQKRKELQENQDEIENMMNSIFKGIFVHRYRDAIAEIRAICIEEIGVWMKMYSDAFLNDSYLKYVGWTLHDRQGEVRLKCLKALQSLYTNRELFPKLELFTNRFKDRIVSMTLDKEYDVAVEAIRLVTLILHGSEEALSNEDCENVYHLVYSAHRPVAVAAGEFLHKKLFSRHDPQAEEALAKRRGRNSPNGNLIRMLVLFFLESELHEHAAYLVDSLWESSQELLKDWECMTELLLEEPVQGEEAMSDRQESALIELMVCTIRQAAEAHPPVGRGTGKRVLTAKERKTQIDDRNKLTEHFIITLPMLLSKYSADAEKVANLLQIPQYFDLEIYSTGRMEKHLDALLKQIKFVVEKHVESDVLEACSKTYSILCSEEYTIQNRVDIARSQLIDEFVDRFNHSVEDLLQEGEEADDDDIYNVLSTLKRLTSFHNAHDLTKWDLFGNCYRLLKTGIEHGAMPEQIVVQALQCSHYSILWQLVKITDGSPSKEDLLVLRKTVKSFLAVCQQCLSNVNTPVKEQAFMLLCDLLMIFSHQLMTGGREGLQPLVFNPDTGLQSELLSFVMDHVFIDQDEENQSMEGDEEDEANKIEALHKRRNLLAAFSKLIIYDIVDMHAAADIFKHYMKYYNDYGDIIKETLSKTRQIDKIQCAKTLILSLQQLFNELVQEQGPNLDRTSAHVSGIKELARRFALTFGLDQIKTREAVATLHKDGIEFAFKYQNQKGQEYPPPNLAFLEVLSEFSSKLLRQDKKTVHSYLEKFLTEQMMERREDVWLPLISYRNSLVTGGEDDRMSVNSGSSSSKTSSVRNKKGRPPLHKKRVEDESLDNTWLNRTDTMIQTPGPLPAPQLTSTVLRENSRPMGDQIQEPESEHGSEPDFLHNPQMQISWLGQPKLEDLNRKDRTGMNYMKVRTGVRHAVRGLMEEDAEPIFEDVMMSSRSQLEDMNEEFEDTMVIDLPPSRNRRERAELRPDFFDSAAIIEDDSGFGMPMFGSGSGGMGDSLSWLLRLLN

<u>SA2</u>

10xHis – SA2 WT – ybbR

MHHHHHHHHHHGSGSSGIAAPEIPTDFNLLQESETHFSSDTDFEDIEGKNQKQGKGKTCKKGKKGPAEKGKGGNGGGKPPSGPNRMNGHHQQNGVENMMLFEVVKMGKSAMQSVVDDWIESYKHDRDIALLDLINFFIQCSGCKGVVTAEMFRHMQNSEIIRKMTEEFDEDSGDYPLTMAGPQWKKFKSSFCEFIGVLVRQCQYSIIYDEYMMDTVISLLTGLSDSQVRAFRHTSTLAAMKLMTALVNVALNLSINMDNTQRQYEAERNKMIGKRANERLELLLQKRKELQENQDEIENMMNAIFKGVFVHRYRDAIAEIRAICIEEIGIWMKMYSDAFLNDSYLKYVGWTMHDKQGEVRLKCLTALQGLYYNKELNSKLELFTSRFKDRIVSMTLDKEYDVAVQAIKLLTLVLQSSEEVLTAEDCENVYHLVYSAHRPVAVAAGEFLYKKLFSRRDPEEDGMMKRRGRQGPNANLVKTLVFFFLESELHEHAAYLVDSMWDCATELLKDWECMNSLLLEEPLSGEEALTDRQESALIEIMLCTIRQAAECHPPVGRGTGKRVLTAKEKKTQLDDRTKITELFAVALPQLLAKYSVDAEKVTNLLQLPQYFDLEIYTTGRLEKHLDALLRQIRNIVEKHTDTDVLEACSKTYHALCNEEFTIFNRVDISRSQLIDELADKFNRLLEDFLQEGEEPDEDDAYQVLSTLKRITAFHNAHDLSKWDLFACNYKLLKTGIENGDMPEQIVIHALQCTHYVILWQLAKITESSSTKEDLLRLKKQMRVFCQICQHYLTNVNTTVKEQAFTILCDILMIFSHQIMSGGRDMLEPLVYTPDSSLQSELLSFILDHVFIEQDDDNNSADGQQEDEASKIEALHKRRNLLAAFCKLIVYTVVEMNTAADIFKQYMKYYNDYGDIIKETMSKTRQIDKIQCAKTLILSLQQLFNEMIQENGYNFDRSSSTFSGIKELARRFALTFGLDQLKTREAIAMLHKDGIEFAFKEPNPQGESHPPLNLAFLDILSEFSSKLLRQDKRTVYVY

LEKFMTFQMSLRREDVWLPLMSYRNSLLAGGDDDTMSVISGISSRGSTVRSKKSKPSTGKRKV
VEGMQLSLTEESSSSDSMWLSREQTLHTPVMMQTPQLTSTIMREPKRLRPEDSFMSVYPMQT
EHHQTPLDYNRRGTSLMEDDEEPIVEDVMMSSEGRIEDLNEGMDFDTMDIDLPPSKNRRERTE
LKPDFFDPASIMDESVLGVSMFGSSGDSLEFIASKLA

Rad21

6x His – Rad21 (Δ1-280; Δ421-631) – TEV site – MBPx

MGSSHHHHHHSGVDPVEPMPTMTDQTTLVPNEEEAFALEPIDITVKETKAKRKRKLIVDSVKEL
DSKTIRAQLSDYSDIVTTLDLAPPTKKLMMWKETGGVEKLFSLPAQPLWNNRLLKLFTRCLTPLV
PEDLRKRRKGGEADNLDEFLKEFGSSGENLYFQGGSGSNSSSGGSGGGSGKIEEGKLVIWING
DKGYNGLAEVGKKFEKDTGIKVTVEHPDKLEEKFPQVAATGDGPDIIFWAHDRFGGYAQSGLLA
EITPDKAFQDKLYPFTWDAVRYNGKLIAYPIAVEALSLIYNKDLLPNPPKTWEEIPALDKELKAKG
KSALMFNLQEPYFTWPLIAADGGYAFKYGDIKDVGVDNAGAKAGLTFLVDLIKNKHMNADTDYSI
AEAAFNKGETAMTINGPWAWSNIDTSKVNYGVTVLPTFKGQPSKPFVGVLSAGINAASPNKELA
KEFLENYLLTDEGLEAVNKDKPLGAVALKSYEEELVKDPRVAATMENAQKGEIMPNIPQMSAFW
YAVRTAVINAASGRQTVDEALKDAQT
